# *Plasmodium falciparum* protein Pfs16 is a target for transmission-blocking antimalarial drug development

**DOI:** 10.1101/2021.06.14.448287

**Authors:** Sabrina Yahiya, Charlie N. Saunders, Ursula Straschil, Oliver J. Fischer, Ainoa Rueda-Zubiaurre, Silvia Haase, Gema Vizcay-Barrena, Sarah Jordan, Sarah Hassan, Michael J. Delves, Edward W. Tate, Anna Barnard, Matthew J. Fuchter, Jake Baum

**Affiliations:** Department of Life Sciences, Imperial College London, Sir Alexander Fleming Building, Exhibition Road, South Kensington, London, SW7 2AZ, UK; Department of Chemistry, Imperial College London, Molecular Sciences Research Hub, White City Campus, Wood Lane, London, W12 OBZ, UK; Centre for Ultrastructural Imaging, New Hunt’s House, Guy’s Campus, King’s College London, London, SE1 1UL, UK

**Author notes:** **Correspondence to:** Jake Baum, Department of Life Sciences, Imperial College London, Exhibition Road, South Kensington, London SW7 2AZ, United Kingdom.

**Keywords:** *Plasmodium falciparum*, malaria, high throughput screening, target identification, target validation, transmission, drug discovery, exflagellation, gametocytes, gametogenesis

## Abstract

Phenotypic cell-based screens are critical to the discovery of new antimalarial lead compounds. However, identification and validation of cellular targets of lead compounds is required following discovery in a phenotypic screen. We recently discovered a *Plasmodium* transmission-blocking N-((4-hydroxychroman-4-yl)methyl)-sulfonamide (N-4HCS) compound, **DDD01035881**, in a phenotypic screen. **DDD01035881** and its potent derivatives have been shown to block *Plasmodium* male gamete formation (microgametogenesis) with nanomolar activity. Here, we synthesised a photoactivatable N-4HCS derivative, probe **2**, to identify the N-4HCS cellular target. Using probe **2** in photo-affinity labelling coupled with mass spectrometry, we identified the 16 kDa *Plasmodium falciparum* parasitophorous vacuole membrane protein Pfs16 as the likely cellular target of the N-4HCS series. Further validating Pfs16 as the cellular target of the N-4HCS series, the Cellular Thermal Shift Assay (CETSA) confirmed DDD01035881 stabilised Pfs16 in lysate from activated mature gametocytes. Additionally, photo-affinity labelling combined with in-gel fluorescence and immunoblot analysis confirmed the N-4HCS series interacted with Pfs16. High-resolution, widefield fluorescence and electron microscopy of N-4HCS-inhibited parasites was found to result in a cell morphology entirely consistent with targeted gene disruption of *Pfs16*. Taken together, these data strongly implicate Pfs16 as the target of **DDD01035881** and establish the N-4HCS scaffold family as a powerful starting point from which future transmission-blocking antimalarials can be developed.

## INTRODUCTION

Malaria continues to devastate millions, with 228 million cases and 405,000 deaths from malaria reported in 2018 alone^1^. The causative agent of malaria, the protozoan *Plasmodium* parasite, transitions between a mammalian host and *Anopheles* mosquito vector, demonstrating extensive cellular plasticity in form across the different lifecycle stages^2^. Symptomatic stages of *Plasmodium* are restricted to the asexual blood stages and can be targeted and rapidly killed by current frontline antimalarials, including artemisinin and its derivatives^3^. Sexual forms (male and female gametocytes) are relatively dormant in the blood and generally more resistant to the effects of conventional antimalarials, yet they are entirely responsible for human to mosquito transmission^4^.

Although huge gains have been made in reducing the burden of malaria since the turn of the millennium, current control measures are threatened by the emergence and spread of parasite resistance to artemisinin-based combination therapies along with mosquito resistance to insecticides^1^. To combat rising parasite resistance, new antimalarial drugs with alternative modes of action are critically needed^3^. Transmission of the parasite from human to mosquito is one of the major bottlenecks in the parasite lifecycle^5^. Given the necessity of transmission to the *Plasmodium* life cycle and proven capacity of transmission-blocking interventions to effectively break the cycle^6^, the process of transmission is emerging as a key target for future antimalarial drug development^7^. A primary target for transmission interventions is the dormant circulating gametocyte, responsible for onwards *Plasmodium* transmission. Drugs that either kill or sterilize these forms in the human host, or prevent fertilisation of resulting gametes in the mosquito gut, would theoretically block transmission^8^. Currently, however, the only approved antimalarials with defined transmission blocking activity are primaquine and tafenoquine of the 8-aminoquinoline family. Use of primaquine is impeded by its clinical safety; the drug being associated with haemolysis in G6PD-deficient patients^9^. New drugs that target the transmission process could therefore have substantial impact in malaria control.

The male and female gametocytes of *P. falciparum* are recognised by their characteristic falciform shape. Gametocytes maintain quiescence as they mature over five distinct morphological stages in the human host, sequestering in the bone marrow and spleen until reaching maturity^10^. Upon reaching maturity, stage V gametocytes re-enter the host circulation where they are available for transmission to *Anopheles* mosquito vectors during a bloodmeal. The formation of gametes from gametocytes, or gametogenesis, occurs in the mosquito midgut. During gametogenesis, male gametocytes transform to 8 haploid male/microgametes (microgametogenesis) whilst female gametocytes form a single haploid female/macrogamete (macrogametogenesis)^11,12^. While females undergo a relatively simple rounding up and erythrocyte egress process, formation of male gametes during microgametogenesis is a notably complex and rapid process. The process involves simultaneous egress from host erythrocytes, three rounds of DNA replication alternating with endomitotic division and eventual formation of axonemes^11,12^. Axonemes, which nucleate from basal bodies, emerge from the parental cell as haploid microgametes in the process of exflagellation. Fertilisation between male and female gametes gives rise to a diploid zygote that, following mosquito midgut colonization, eventually yields haploid motile sporozoites responsible for the next transmission cycle to humans^2^.

Efforts to identify novel drugs targeting processes from symptomatic blood through to transmission have been significantly bolstered by advances in high throughput phenotypic screening of compound libraries^7^. However, whilst phenotypic screens supersede target-based drug discovery in their ability to rapidly identify hits, the lack of knowledge on the hit target or its mode of action can result a lengthy process of development or safety concerns during clinical testing. Elucidation of a hit compound’s cellular target(s) and mode of action prior to clinical testing is therefore essential for improving the success rates of drug candidates^13^. One recent exemplar high throughput screen involved testing of a large diverse chemical library (the Global Health Chemical Diversity Library (GHCDL)) against both transmissible and symptomatic stages of *Plasmodium*. The GHCDL screen identified hits with diverse activity profiles against the *Plasmodium* lifecycle, including hits with multistage, transmission-specific, sex-specific and stage-specific activity^14^. Within the library, a series of compounds with an N-((4-hydroxychroman-4-yl)methyl)-sulfonamide (N-4HCS) scaffold were found to potently inhibit formation of *Plasmodium* male gametes from mature stage V male gametocytes. The most potent hit from this scaffold series, **DDD01035881**, has since been extensively studied by structure activity relationship, yielding hits with half maximal inhibitory (IC_50_) concentrations in the nanomolar range^15^. Importantly, **DDD01035881** and derivatives were also shown to have minimal impact on viability of HepG2 mammalian cells, suggesting they have low toxicity^15^.

To advance the N-4HCS scaffold here we sought to identify the mode of action and/or cellular target of **DDD01035881**. Using photo affinity labelling, label-free Cellular Thermal Shift Assay (CETSA) and cellular analysis of treated parasites, we present strong interdisciplinary evidence that the target of **DDD01035881** and N-4HCS scaffold is the *Plasmodium falciparum* 16 kDa parasitophorous vacuole membrane protein, Pfs16. Given the power of transmission blocking therapeutics and drive for discovery of further, novel combination therapies, these data suggest the N-4HCS scaffold may be an excellent foundation for future antimalarial treatment or preventative regimens and positions Pfs16 as a novel target for further development.

## RESULTS

The N-4HCS scaffold was recently identified in a high-throughput screen for transmission blocking antimalarials^14^, with subsequent development to improve its activity by medicinal chemistry^15^. We sought to determine a potential mode of action and cellular target for the N-4HCS scaffold, starting out from the original hit **DDD01035881**, given its potency for inhibiting *Plasmodium falciparum* parasite microgametogenesis, the process of male gamete formation from blood-circulating mature male gametocytes^14^.

### Identification of gametocyte-specific targets of the N-4HCS scaffold

To facilitate identification of potential cellular target(s) via photoaffinity labelling we first derivatised **DDD01035881** to incorporate both a photo-activatable group and clickable alkyne moiety onto the N-4HCS scaffold (see **Supplementary Information** for chemistry). Despite a number of designs, the tolerance of the N-4HCS scaffold to large changes in structure was found to be limited (**Table S1**), consistent with our previous medicinal chemistry study^15^. Ultimately, a strategy to incorporate an alkyne handle and aryl azide moiety separately on the molecule was developed, leading to the synthesis of probe **2**. Parent molecule **1** was synthesised to resemble probe **2** by retaining similar structural changes to the N-4HCS scaffold without the photoactivatable and clickable moieties, thus mimicking biological activity of probe **2** as a control or active competitor. Critically, parent molecule **1** and probe **2** retained micromolar to nanomolar IC_50_s in the *in vitro* male gamete formation assay, respectively (**Table 1**).

**Table 1.**
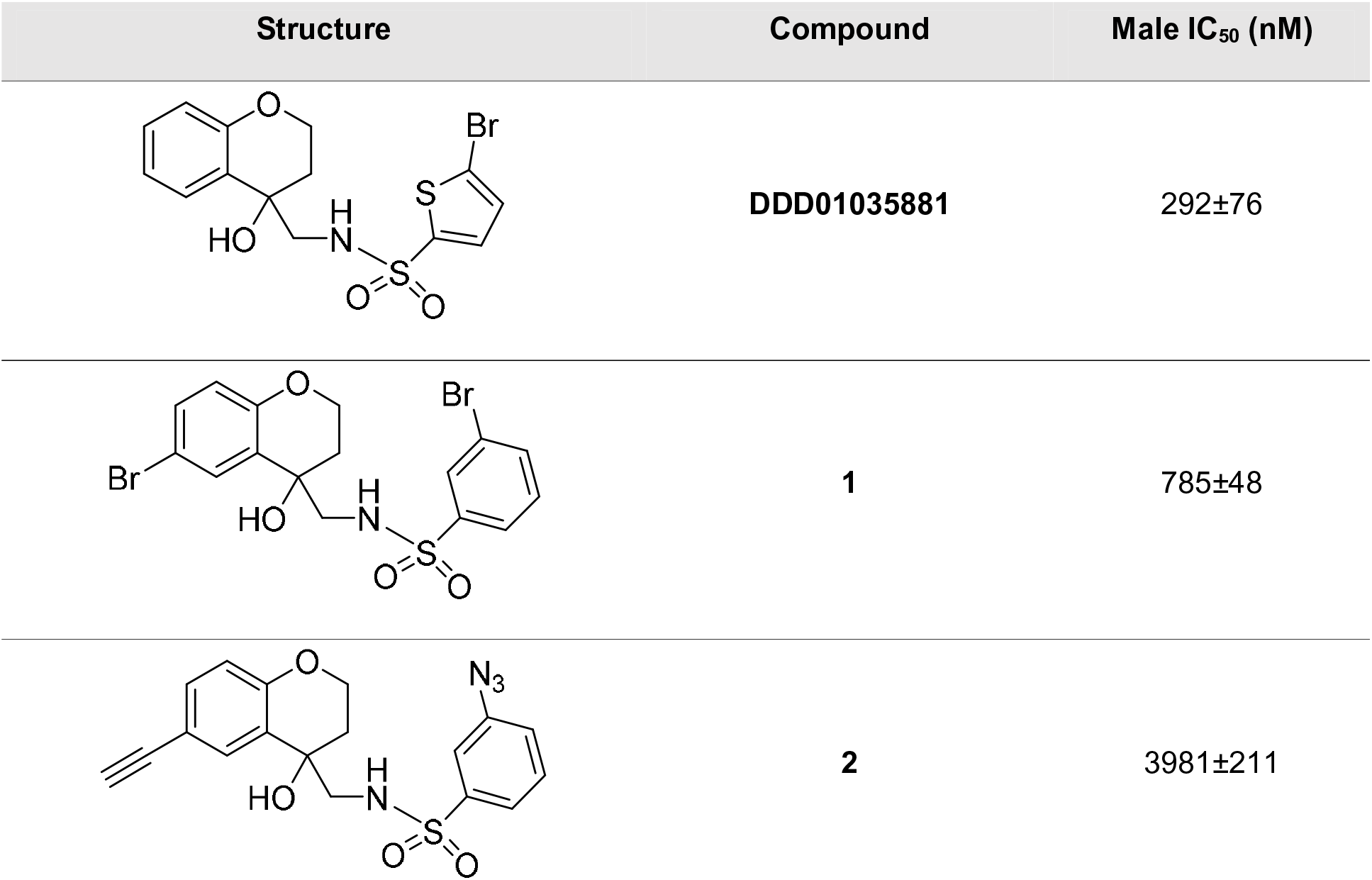
DDD01035881 and derivatives. Activity of **DDD01035881** and structural analogues as determined in a structure activity relationship study. IC_50_s depicted represent the inhibition of male gamete viability as determined by the male gamete formation format of the DGFA. Values are expressed as an average ± SEM of 3-8 biological replicates, ≥ 2 technical replicates.^15^

| Structure | Compound | Male IC<sub>50</sub> (nM) |
| --- | --- | --- |
|  | **DDD01035881** | 292±76 |
|  | **1** | 785±48 |
|  | **2** | 3981±211 |

Preliminary testing of probe **2** was performed to test cross-linking and ligation to azido-TAMRA/biotin capture reagent (AzTB)^16^. Cell lysate derived from activated mature gametocytes was irradiated in the presence of increasing concentrations of probe **2** or DMSO to test a probe **2** concentration-dependent crosslinking of proteins. Lysate containing crosslinked proteins was then ligated to AzTB capture reagent in a copper catalysed azide-alkyne cycloaddition (CuAAC). Crosslinked and AzTB-ligated proteins were then enriched with streptavidin coated beads. Protein labelling of enriched proteins, as determined by in-gel fluorescence (IGF), was found in the presence of probe **2** and was additionally shown to be AzTB dependent, thus confirming successful crosslinking and CuAAC click reactions (**Figure S1A**).

Target identification was subsequently carried out using a 9plex tandem mass tag (TMT) methodology. Live *P. falciparum* stage V gametocyte cultures were treated with either DMSO, probe **2** (10 µM) or a combination of probe **2** and parent molecule **1** to act as a competitor (10 µM each), acquiring triplicate samples of each condition. Live treated gametocytes were irradiated at 254nm to crosslink probe to protein targets, then purified and lysed before ligating to AzTB in a CuAAC reaction. AzTB-ligated proteins were subsequently enriched with Neutravidin agarose beads before preparing peptides for TMT labelling and quantification. Peptide samples were then prepared for analysis by nanoscale liquid chromatography mass spectrometry on a QExactive orbitrap mass spectrometer (nLC-MS/MS). Analysis of peptides found by mass spectrometry revealed 129 protein hits (for raw data see **Supporting Information**).

Of 129 protein hits identified, the specific probe-protein interaction profile was determined by omitting any hits identified in DMSO-treated samples, excluded as non-specific binding interactions. Comparing DMSO and probe **2** treated fractions, probe-specific interactions revealed 20 protein hits positively enriched as a specific result of probe **2** labelling and AzTB ligation, as shown in **Figure 1A**. To further confirm the specificity of the interaction between probe and protein hits, samples treated with probe **2** in the presence of parent molecule **1**, serving as a competitor, were compared to samples treated with probe **2** alone. Compared in this way, out-competed probe-specific interactions revealed 9 protein hits, depicted as negatively enriched proteins in **Figure 1B**. Of these 9 hits, 6 were also positively enriched in the initial comparison of DMSO and probe **2**-treated fractions (**Figure 1A**, **Table 2** and see also **Extended SI** and **Figure S1B-C**).

**Figure 1.**
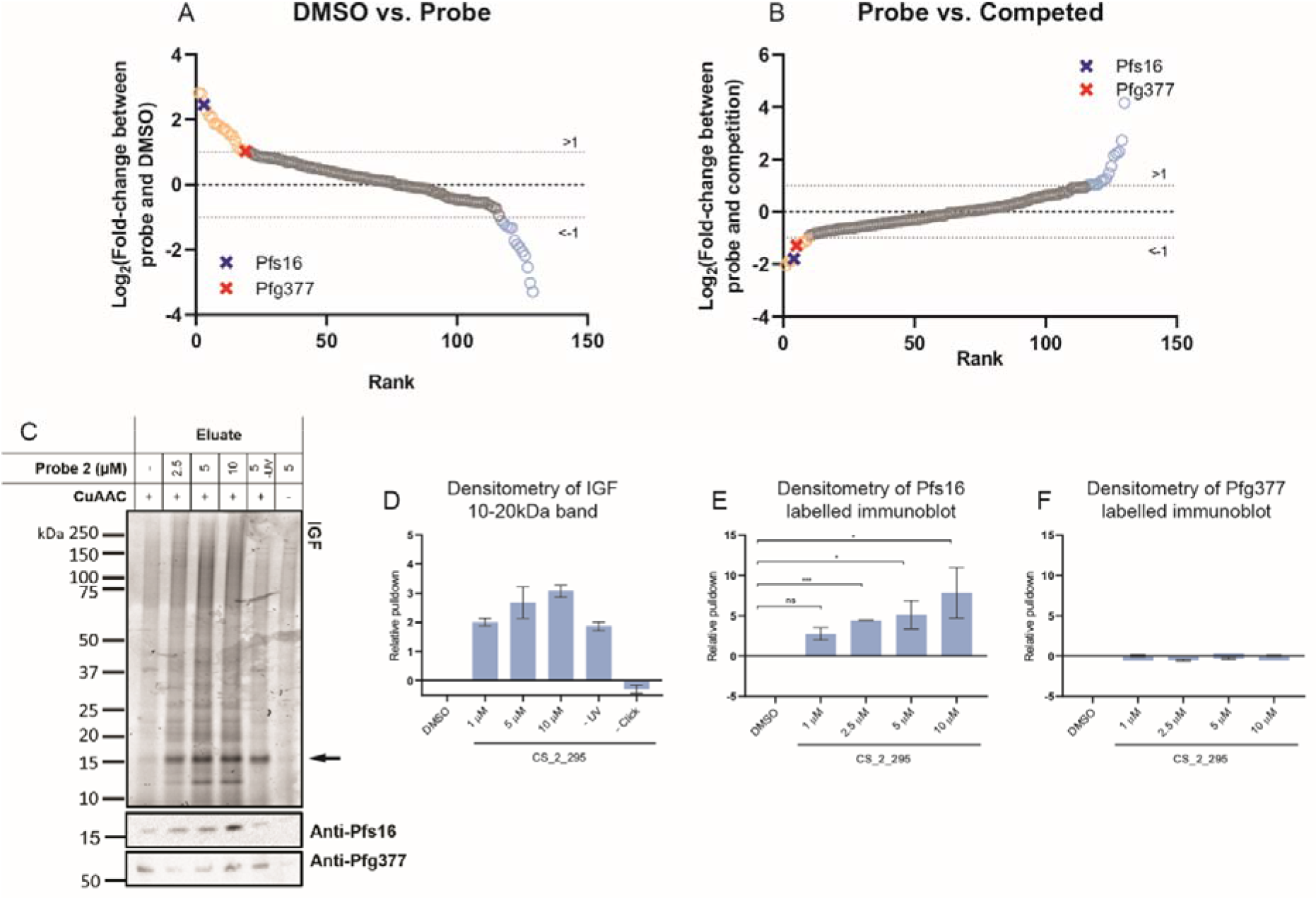
Identification and validation of Pfs16 as a specific interaction partner with the N-4HCS series. ^61^ Results of *P. falciparum* proteome-wide target identification study. Proteins enriched by PAL and pulldown were identified by nLC-MS/MS. Plots depict the log_2_-transformed fold change in protein enrichment between the of **(A)** DMSO and probe **2** treated samples or **(B)** probe **2** and competition (combined treatment probe **2** and parent molecule **1**) samples. All protein hits, represented as circles, are ranked based on the log_2_-transformed fold change in enrichment, between the average values of samples. Averages were taken of 3 distinct biological replicates. Orange circles denote positive log-transformed fold change >1 and blue circles denote negative log-transformed fold change <-1 in **A**, and vice-versa in **B**. Pfs16 and Pfg377 are marked with blue and red circles, respectively, in both **A** and **B**. Hits enriched in both conditions are listed in **Table 2**. **(C-F)** Validation of Pfs16 and Pfg377 binding by live probe **2** treatment, AzTB conjugation, streptavidin-biotin affinity enrichment and analysis by IGF and immunoblot. IGF fluorescence was used to identify streptavidin-enriched proteins and corresponding immunoblots validated the specificity of pulldowns to Pfs16 and Pfg377. **(C)** IGF revealed an abundantly TAMRA-labelled protein between 15kDa and 20kDa in size, likely corresponding to Pfs16. Band intensity, relative to DMSO, is depicted as an average and error bars denote SEM of 3 biological replicates. See **Figure S2B-C** for further replicates. **(D)** Densitometry of the 16kDa protein band in **C** and **Figure S2B-C** was determined and is depicted as relative band intensity relative to a DMSO control in the presence of increasing probe **2** concentration. Values depict averages and error bars denote SEM of 3 biological replicates. Densitometry of corresponding immunoblots labelled with **(E)** Pfs16 and **(F)** Pfg377 are depicted as band intensities relative to a DMSO control, in the presence of increasing probe **2** concentration. Error bars denote SEM of 3 biological replicates, see **Figure S2** for immunoblots and gels which were analysed to obtain **(D-F)**. Significance was determined by performing an unpaired two-tailed t-test and is denoted as *** (p < 0.001), * (p < 0.05) and ns (p ≥ 0.05).

**Table 2.** List of protein hits enriched by PAL and nLC-MS/MS. Protein hits identified as specific to the probe **2**-protein interaction profile, including *Pfs16 and **Pfg377, ranked according to the log-transformed fold change (Log_2_FC) in protein enrichment between average values of 3 biological replicates. Proteins depicted were identified as both positively enriched when comparing DMSO and probe 2, and, negatively enriched when comparing probe 2 and a combination of probe 2 and parent molecule 1. P values listed are derived from analysis by student’s *t*-test, see **Supporting Information**.

| Rank | DMSO vs. probe 2 |  |  |  | Probe 2 vs. competition (probe 2 + parent molecule 1) |  |  |  |
| --- | --- | --- | --- | --- | --- | --- | --- | --- |
|  | Colloquial Name | Protein ID | Log<sub>2</sub>FC | P value | Colloquial Name | Protein ID | Log<sub>2</sub>FC | P value |
| 1 | **\*Pfs16** | **PF3D7_0406200.1-p1** | 2.45809 | **0.0701** | Spermidine Synthase | PF3D7_1129000.1-p1 | -2.0397 | 0.1552 |
| 2 | Spermidine Synthase | PF3D7_1129000.1-p1 | 2.27369 | 0.1058 | 60S Ribosomal Protein L26 | PF3D7_0312800.1-p1 | -1.8946 | 0.0051 |
| 3 | RAB-1B (Ras related protein) | PF3D7_0512600.1-p1 | 2.08532 | 0.0314 | RAB-1B (Ras related protein) | PF3D7_0512600.1-p1 | -1.8675 | 0.0177 |
| 4 | 60S Ribosomal Protein L26 | PF3D7_0312800.1-p1 | 1.71633 | 0.0423 | **\*Pfs16** | **PF3D7_0406200.1-p1** | **-1.7926** | **0.3069** |
| 5 | V-type Proton ATPase | PF3D7_1311900.1-p1 | 1.34692 | 0.0567 | **\*\*Pfg377** | **PF3D7_1250100.1-p1** | **-1.2868** | **0.1180** |
| 6 | **\*\*Pfg377** | **PF3D7_1250100.1-p1** | 1.02634 | **0.4615** | V-type Proton ATPase | PF3D7_1311900.1-p1 | -1.1617 | 0.0201 |

Among the 6 protein hits specific to probe **2** (**Table 2**), the protein most positively enriched when comparing DMSO-treated and probe **2**-treated samples was the gametocyte-specific 16 kDa *Plasmodium falciparum* vacuole membrane protein Pfs16 (PlasmoDB ID, PF3D7_0406200)^17^. The *Plasmodium falciparum* female gametocyte-specific protein Pfg377 (PlasmoDB ID, PF3D7_1250100)^18^ was also identified. As the N-4HCS compounds have been shown to solely inhibit gametocyte viability but to exert no effect on other *Plasmodium* lifecycle stages^14^, Pfs16 and Pfg377 were prioritised for further study. All other protein hits identified here were not specific to the gametocyte stages and were therefore deemed likely to be the result of non-specific binding interactions. For example, spermidine synthase, a well characterised enzyme is expressed specifically during erythrocytic schizogony^19^ and inhibitors of the enzyme have been shown to impair asexual growth^20,21^, making it an unlikely target of the gametocyte-specific N-4HCS series. Rab1B, is believed to have a potential role in ER to Golgi transport^22^ and shown to lie adjacent to the ER in early asexual blood stages^23^. Since the *Rab1B* gene is essential for asexual blood stage parasite growth is it also likely the result of non-specific binding^24^. The V-type proton ATPase has been shown to function in the asexual blood stages^25^ where it plays a critical role in maintaining the parasite cytosol pH^26^. Specific inhibitors of V-type proton ATPase have been shown to markedly impact asexual growth^27,28^, making it an unlikely target of the N-4HCS series. Much like any ribosomal targeted protein, including the hit L26, it is unlikely that the transmission specific effects of the N-4HCS scaffold would be mediated via inhibition of these proteins. Thus, we focused our attention on the sexual-stage specificity of the N-4HCS series, prioritising hits Pfs16 and Pfg377 for further validation.

Pfs16 is a 157-amino acid protein with two transmembrane domains and is one of the earliest known markers of sexual conversion in *Plasmodium*^29^. Importantly, with respect to validating the phenotype described for the N-4HCS scaffold^14^, knockout of the *Pfs16* gene is known to be associated with a block in microgametogenesis as is also evident with N-4HCS compound treatment ^30^. Pfg377 is, in contrast, only associated with female gametocytes and would therefore be less likely as a target of the N-4HCS scaffold which has been shown to specifically inhibit male gamete formation. Given its phenotypic consistency with drug action, we therefore sought to further validate Pfs16 as a cellular target of the compound whilst simultaneously investigating whether the Pfg377 interaction is specific.

### Investigation of Pfs16 and Pfg377 as potential targets of the N-4HCS scaffold by PAL

To validate TMT-dependent identification of Pfs16 and Pfg377 as potential N-4HCS targets, PAL was repeated and analysed by IGF and immunoblot. As with the TMT identification, live mature gametocytes were treated with increasing concentrations of probe **2** and irradiated with UV to enable bioconjugation to cellular target(s). The probe-tagged proteins present in the resulting cell lysate were biotinylated with TAMRA-containing AzTB capture reagent^31^ and pulled down with streptavidin coated magnetic beads. Enriched proteins which were ligated to AzTB and thus TAMRA-labelled were then analysed by IGF, performing the experiment in triplicate. A notable protein band between 15 and 20kDa, likely corresponding to the 16kDa protein Pfs16, demonstrated a probe **2** concentration-dependent pulldown relative to DMSO in each replicate (**Figure 1C**). Densitometry revealed that the relative band intensity of the enriched protein band increased with higher concentrations of probe **2** relative to a DMSO control, over each replicate (**Figure 1D**). Corresponding immunoblot and densitometry was then used to analyse the specificity of the pulldown to Pfs16 and Pfg377, by using increasing concentrations of probe **2**, and antibodies specific to each of the proteins (see **Figure S2** for the full triplicate IGF data of protein pulldowns and corresponding immunoblots). While a dose-dependent pulldown of Pfs16 was clearly observed (**Figure 1E**), there was no correlation between probe **2** concentration and the amount of Pfg377 pulled down by the probe (**Figure 1F**), supporting our hypothesis of the latter being a non-specific interaction. The dose-dependent pulldown of Pfs16, in contrast, adds further support to Pfs16 being the likely target of the N-4HCS series.

Adding further validation, pulldowns were performed on parasites treated with a combination of probe **2** and parent compound **DDD01035881**, acting as a competitor. If the pulled-down protein is a target of the N-4HCS scaffold, addition of a highly potent competitor should result in a decrease in the amount of protein available for pull-down by the probe. A clear decrease in the protein band detected at 15-20 kDa by IGF was indeed seen when comparing the pulldowns in the presence and absence of parent molecule (**Figure S3**). This result confirms specific pull down of Pfs16 following probe **2** labelling, attributable to the N-4HCS scaffold and hence to the **DDD01035881** series.

### Label-free validation of Pfs16 as a potential target of DDD01035881

PAL is known to be vulnerable to false positive results, with clickable derivatives potentially binding non-specifically to proteins besides the target of interest^32^. To validate the specificity of **DDD01035881** engagement with proteins Pfs16 and Pfg377, the label-free Cellular Thermal Shift Assay (CETSA) was used with lysate from mature gametocyte cultures and the original **DDD01035881** compound. CETSA is based on the premise that proteins irreversibly aggregate when thermally challenged and the modulation of a given protein when bound to a ligand can alter this process, resulting in increased thermal stability^33^. Using a thermal gradient, a melting curve of a given protein can then be obtained to compare the protein’s melting temperature (T_m_) and the temperature at which the protein aggregates in the presence and absence of a ligand^34^. A positive shift in a protein’s T_m_ relative to a DMSO control would indicate protein stabilisation due to ligand engagement, confirming drug binding to the protein. Melting curves and T_m_ can be obtained by analysis of corresponding immunoblots using antibodies specific to the protein of interest, permitting comparison of T_m_ as relative immunoblot density. We utilised in-lysate CETSA using 1% Triton-X100-containing lysis buffer, which would be expected to solubilise a PVM protein target such as Pfs16, prior to compound treatment and thermal challenge. The lysis conditions used matched those used in the PAL target identification study.

Mature gametocyte cell lysate from activated stage V gametocytes was incubated with either **DDD01035881** or DMSO as a control, thermally challenged and probed by immunoblot using Pfs16-specific antibody^17^ to explore the engagement of **DDD01035881** and Pfs16 (**Figure S4**). Pfs16 melt curves were obtained to compare the T_m_ between DMSO-treated and **DDD01035881**-treated soluble gametocyte protein fractions (**Figure 2**). **DDD01035881** treatment clearly resulted in a positive shift in the Pfs16 melting curve (**Figure 2A**). When compared to DMSO by densitometry, the relative band-density of **DDD01035881** treated fractions were found to be significantly different at 91°C (unpaired two tailed t test, p < 0.001). Utilising the same approach, **DDD01035881** did not show significant stabilisation of Pfg377 at any temperature investigated (**Figure 2B**). This aligns with the published female-specific role of the Pfg377 in macrogametogenesis and lack of activity of **DDD01035881** against female gametocytes^18^. These results strongly support a specific interaction between **DDD01035881** and Pfs16 in mature activated male gametocytes, corroborating the PAL. The lack of interaction with the female-specific Pfg377 suggests its detection by PAL was a false-positive, consistent with our understanding that **DDD01035881** specifically targets males.

**Figure 2.**
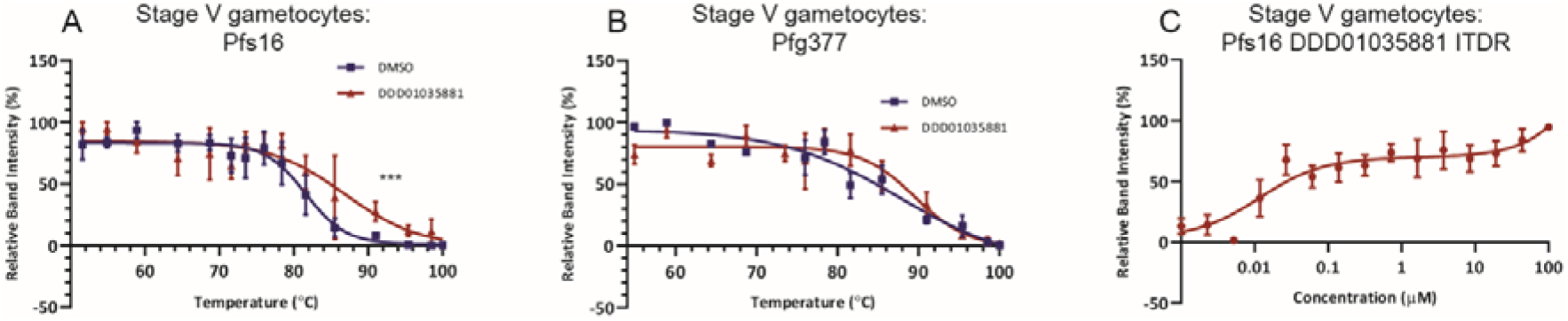
Label-free validation of Pfs16 interaction with DDD01035881 in mature gametocytes. **(A-C)** Target validation by CETSA, using densitometric analysis of immunoblots for melt curve generation and subsequent quantification of Tm. Melt curves demonstrating thermal stability of **(A)** Pfs16 and **(B)** Pfg377 using activated stage V gametocyte lysate treated with DMSO or **DDD01035881**. Error bars represent the SEM of 2-3 biological replicates. In **(A)**, a statistically significant difference was found at 91°C using an unpaired two-tailed t-test (p < 0.001). **(C)** The corresponding isothermal dose response (ITDR) curve of **(A)** depicting the concentration dependent stabilisation of Pfs16 by **DDD01035881** in activated stage V gametocyte lysate. Error bars represent the SEM of 4 biological replicates. Full immunoblots can be found in **Figure S4**.

To further validate the positive-shift in Pfs16 T_m_ with **DDD01035881** treatment of activated gametocyte lysate, an isothermal dose response (ITDR) format of CETSA was applied. ITDR-CETSA utilises the same premise as the melt curve format, but proteins are thermally challenged with a single temperature and treated with varying compound concentrations^35^. The single temperature applied in ITDR-CETSA is that at which Pfs16 is mostly aggregated in the DMSO-treated fraction but is largely stabilised in the **DDD01035881** fraction (derived from the melt curve, **Figure 2A**). ITDR-CETSA with **DDD01035881** was performed at 78.4°C using concentrations between 1 nM and 100 µM with the results analysed by immunoblotting (**Figure 2C**). A clear concentration dependent stabilisation of Pfs16 was seen, adding substantial support to Pfs16 being a specific interactor of the label-free **DDD01035881**.

### DDD01035881 specifically inhibits microgametogenesis without impacting gametocytogenesis

Pfs16 is reported to be the earliest marker of sexual conversion, with gene disruption suggesting it plays a crucial role in commitment to gametocytogenesis^30^. We therefore sought to study the effect of **DDD01035881** treatment on sexual conversion and early gametocyte development. To determine the stage specificity of Pfs16 binding, CETSA was performed on immature gametocyte cell lysate derived from stage I-III gametocyte culture. Following an identical experimental approach, stage I-III gametocyte lysates were treated with **DDD01035881** and thermally challenged to quantify the stabilisation of Pfs16 and Pfg377. No positive shift or statistically significant difference was found between the DMSO and **DDD01035881** treated fractions in both Pfs16 (**Figure 3A**) and Pfg377 (**Figure 3B**). These findings suggest that binding of **DDD01035881** is specific to Pfs16 in mature gametocytes, with no binding observed for Pfs16 in immature gametocytes or Pfg377 at any stage.

**Figure 3.**
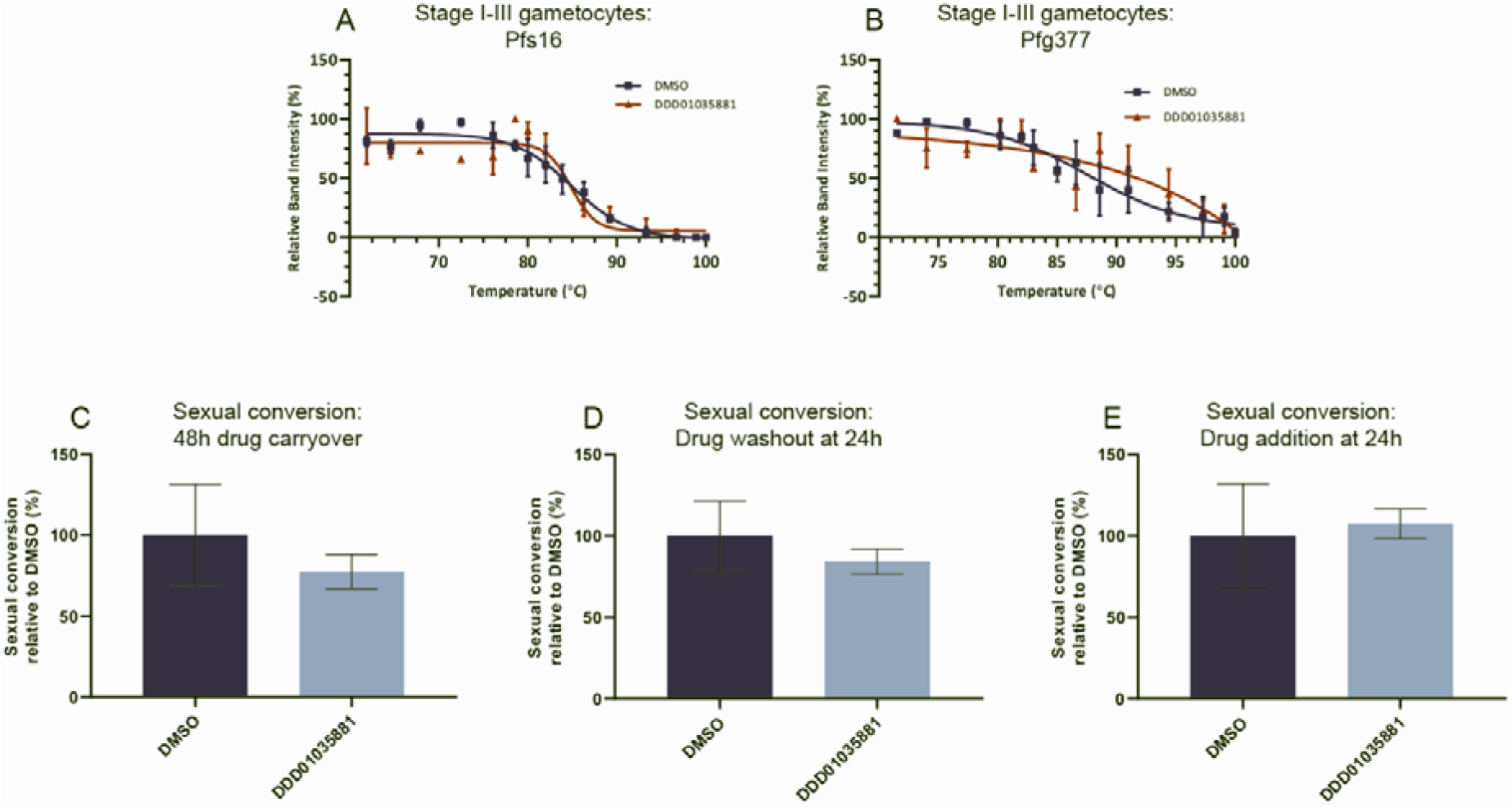
Effect of N-4HCS series on gametocytogenesis. Elucidation of the effect of the N-4HCS series on immature gametocytes by CETSA and **(C-E)** sexual conversion rate calculations. Label-free validation of the effect of **DDD01035881** treatment to sexual conversion by CETSA was performed on lysate derived from stage I-III gametocytes, looking specifically at the stabilisation of **(A)** Pfs16 and **(B)** Pfg377. No statistically significant stabilisation of either Pfs16 or Pfg377 was found at any given temperature between DMSO and **DDD01035881**. Error bars represent the SEM of 2-3 biological replicates. Full immunoblots can be found in **Figure S4**. **(C-E)** The sexual conversion rate of Pf2004/164-tdTomato parasites was determined by quantification of tdTomato expression after induction of gametocytogenesis. Conversion rates of **DDD01035881**-treated parasites are expressed as rates relative to the conversion rates of DMSO-treated parasites. Perturbations to sexual conversion were determined by **(F)** maintaining treatment over two intraerythrocytic cycles and **(G)** the reversibility of any perturbations were determined by removing compound at 24 hours. **(H)** Perturbations to early gametocyte development were probed by administration of compounds in a subsequent intraerythrocytic cycle. Error bars represent the SEM of 2 biological replicates. See **Figure S5** for the gating strategy and conversion rates of further N-4HCS analogues.

Validation of the stage specificity of Pfs16 binding *in vitro* was determined by measuring conversion rates of a transgenic *P. falciparum* line, which expresses tdTomato at the point of sexual conversion (*Pf*2004/164-tdTomato^36^) as determined by flow cytometry (**Figure S5**). Conversion rates of parasites treated with 4-NHCS compounds were quantified relative to a negative control. Here, gametocytes were treated with either **DDD01035881** or a DMSO under multiple conditions and used to quantify i) inhibition of sexual conversion (**Figure 3C**), ii) the reversibility of any inhibitory effect on conversion (**Figure 3D**) and finally, iii) effects to early gametocyte development (**Figure 3E**). The respective conditions were i) prolonged compound exposure from the point of induction (**Figure 3C**), ii) compound removal 24 hours post-induction (**Figure 3D**) and iii) late compound treatment 24 hours after induction (**Figure 3E**, the gating strategy and conversion rates of additional N-4HCS compounds can be found in **Figure S5**). We found no significant reduction in conversion rates under any of the three treatment conditions, suggesting that **DDD01035881** action functions specifically during microgametogenesis and not sexual conversion or early gametocyte development.

### DDD01035881 activity window coincides with Pfs16 activity during microgametogenesis

Having defined that the N-4HCS compound series likely targets Pfs16 during microgametogenesis, we next sought to assess the precise cellular phenotype of the parent molecule, **DDD01035881**, beginning with defining its window of action within the process of microgametogenesis. As **DDD01035881** is known to inhibit microgametogenesis without requiring prior incubation with gametocytes, we hypothesised the compound may continue to exert inhibitory activity beyond activation of gametocytes. To narrow down an activity window for N-4HCS compounds, gametocytes were activated in the absence of **DDD01035881** and subsequently treated with **DDD01035881** in time increments up to the point of exflagellation. Exflagellation rates were counted at 25 mins post-activation and calculated as a percentage relative to DMSO (i.e. untreated) controls. As depicted in **Figure 4A**, **DDD01035881** inhibited microgametogenesis up to 6 min after gametocyte activation. This result is consistent with the time window when the parasitophorous vacuole membrane (PVM), the membrane in which Pfs16 is found, is known to function in microgametogenesis^37^.

**Figure 4.**
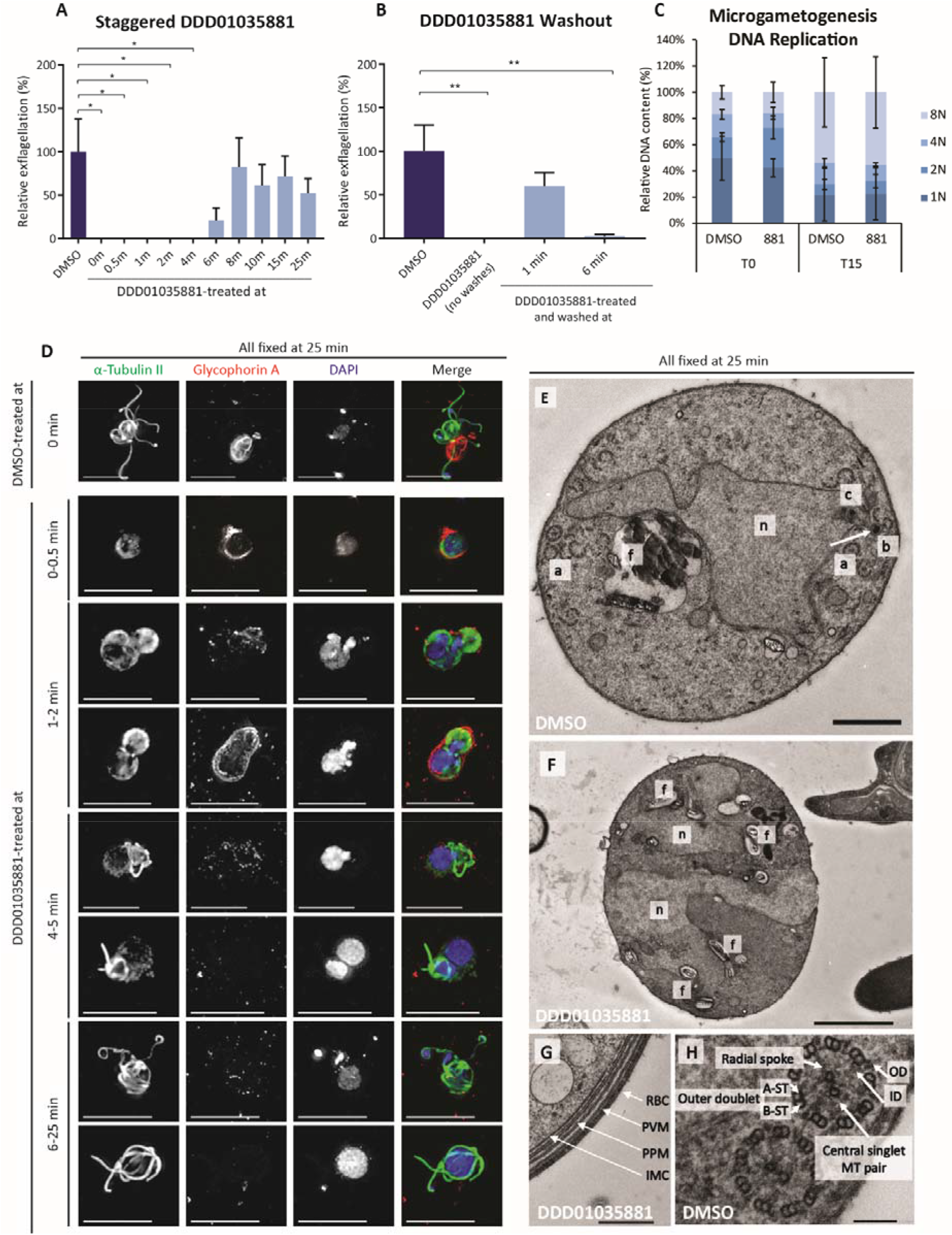
Microgametogenesis phenotype under DDD01035881 treatment. Exflagellation rates of gametocytes relative to DMSO controls which were either **(A)** activated in the absence of drug, then treated with 5µM **DDD01035881** at the stated time points or **(B)** treated with **DDD01035881** or DMSO, then activated and washed three times to remove compound at the labelled time-points (or left under **DDD01035881** pressure without washing). Exflagellation was counted at 25 minutes and error bars represent the SEM of 5-6 biological replicates. Significance was determined with an unpaired two-tailed t-test and is denoted as * (p < 0.05) and ** (p < 0.01). **(C)** Relative DNA content of *PfDyn*GFP/*P47*mCherry male gametocytes treated with either DMSO or **DDD01035881** (denoted by 881) at 0 min and measured at 0 and 15 minutes post-activation. Ploidy of parasites (1n, 2n, 4n or 8n) was determined by flow cytometry as a measure of Vybrant™ DyeCycle™ intensity. Error bars represent the SEM of 3 biological replicates. **(D-H)** Morphological phenotype of **DDD01035881** treated parasites as determined by fluorescence and electron microscopy. **(D)** IFAs of abhorrent DDD0103588-treated male gametocytes, depicting alpha tubulin-labelled cytoskeleton (green), glycophorin A-labelled erythrocyte (red) and DNA (blue). Parasites were treated at the stated time points relative to activation of microgametogenesis and fixed at 25 minutes post-activation, the timepoint at which DMSO treated control parasites exflagellate. Scale bars = 10µm. **(E-H)** EM images of gametocytes treated with **DDD01035881** or DMSO and then activated and fixed at 25 minutes. **(E)** A DMSO-treated male gamete preparing for emergence from the gametocyte cell body. The kinetosomal sphere and granule and kinetosomal basket (b) located at the centriolar plaque within the nucleus (n) which bears an intranuclear spindle (s) and chromatin (c). The food vacuole (f) near to the nucleus. Both normal and aberrant 9+2 axoneme arrangement (a) at the periphery of the cell, preparing for axoneme emergence from the cell body. Scale bar = 1µm. **(F) DDD01035881**-treated microgametocyte with a disturbed nuclear structure (n) and food vacuoles (f). Scale bar = 2µm. **(G) DDD01035881**-treated microgametocyte which has failed egress from the host erythrocyte, displaying an intact 4-layer membrane. Scale bar = 500nm. **(H)** The characteristic 9+2 arrangement of microtubules. A and B subtubule pairs are spaced apart from the central singlet microtubule (MT) pair using radial spokes. Scale bar = **100nm**.

To determine whether **DDD01035881** treatment is reversible during the active window, gametocytes were next treated with **DDD01035881** at the point of activation, washed to remove compound and exflagellation rates measured. Reversibility of **DDD01035881** was shown to be time dependent. Removal of the compound at 1 min restored exflagellation, however, inhibition was retained when the compound was removed at 6 min (**Figure 4B**), the point at which **DDD01035881** loses activity. Combining these observations, it can clearly be concluded that inhibition by **DDD01035881** reversibly blocks microgametogenesis within a 6 min activity window post-gametocyte activation. Beyond 6 min post-activation, the Pfs16-containing PVM ^17,29,38^, is lost due to microgametocyte egress^37^ and hence, loss of **DDD01035881** activity beyond the 6 min window points to reversible binding to Pfs16. It is likely that upon reversibly binding to Pfs16, **DDD01035881** results in the downstream inhibition of exflagellation which points to a corresponding window of Pfs16 activity.

### DDD01035881 treatment does not impact ploidy during microgametogenesis

We next sought to decipher whether **DDD01035881** treatment plays a role in DNA replication, one of the key events in microgametogenesis. To determine the ploidy of gametocytes we used flow cytometry analysis of a transgenic parasite, *PfDyn*GFP/*P47*mCherry, that expresses GFP in male or mCherry in female gametocytes (**Figure S6A**)^39^. Vybrant™ DyeCycle™ Violet staining was used as a measure of male gametocyte DNA content at 0 and 15 min post-activation. To measure ploidy, the GFP and Vybrant™ DyeCycle™ Violet double-positive gametocyte population was gated, from which discrete populations of 1n, 2n, 4n and 8n male gametocytes could then be measured (**Figure S6B-D**).

For DMSO treated control parasites, at 0 min, gametocytes with 1n genome were the most abundant, with a smaller proportion having 2n, 4n or 8n genomes. This small proportion of cells with evidence of genome replication, is a probable result of premature activation or selective gene amplification^40^. Conversely, most gametocytes had an 8n genome at 15 min post-activation, which is indicative of three successful rounds of DNA replication^40^. A reduced proportion of gametocytes failed to fully replicate DNA, with all 1n, 2n and 4n genomes found at different ratios. Ploidy of **DDD01035881**-treated and DMSO-treated gametocytes were found to be similar at both 0 and 15 min post-activation, with no statistically significant difference found (**Figure 4C**). Thus, treatmen with **DDD01035881** does not lead to a defect to in replication during microgametogenesis. Thus, if **DDD01035881** targets Pfs16, by extension this suggest that Pfs16 does not function in or signal upstream of DNA replication (**Figure 4C**).

### DDD01035881 treatment disrupts cytoskeletal, nuclear and food vacuole structure

Having defined that **DDD01035881** activity is specific to microgametogenesis without impacting gametocytogenesis, we next sought to define the compound phenotype during microgametogenesis. Perturbances to microgametogenesis under **DDD01035881** treatment were analysed by either immunofluorescence (IF) microscopy or electron microscopy, using DMSO-treated gametocytes as a control (**Figure S7**).

First, IF analysis was used to determine the effects of compound treatment on cytoskeletal rearrangement, host erythrocyte egress and DNA replication. Gametocytes were first activated in the absence of drug and then treated with **DDD01035881** at various timepoints relative to activation, before being fixed and stained for analysis. As shown in **Figure 4D**, **DDD01035881** treatment resulted in a perturbance to microgametogenesis compared to DMSO-treated gametocytes. The morphological phenotype found under **DDD01035881** treatment was found to be treatment-time dependent with distinct phenotypes observed depending on time of treatment relative to activation.

**DDD01035881** treatment at 0-0.5 min post-activation blocked exflagellation and the cytoskeletal rearrangement of parasites, with gametocytes failing to form mitotic spindles or axonemes. Gametocytes succeeded in rounding up, but the egress phenotype was mixed, with some parasites failing and some succeeding to egress from the host erythrocyte (**Figure 4D**). From 1-2 min, exflagellation failed and the parasite cytoskeleton, depicted as labelled alpha tubulin, adopted a figure-of-eight morphology. The mixed egress phenotype was retained, with erythrocyte vesicles remaining close to the parasite as expected in microgametogenesis^37^; those failing to egress also demonstrated some erythrocyte vesiculation, although to a lesser extent. DNA staining suggested replication was successful, with DNA either localising to one or both sides of the figure-of-eight. Similarly, treatment from 4-5 min resulted in a cytoskeletal figure-of-eight morphology, but egress and erythrocyte vesiculation was successful. However, a truncated flagellar formed from the larger side of the figure-of-eight with tubulin staining markedly more diffuse on the opposing end of the parasite. From 6-25 min, exflagellation appeared to match that of the DMSO control (**Figure 4D**), although the onwards viability of gametes was not determined. Should Pfs16 be the target of **DDD01035881**, this points to a potential function of the protein upstream of cytoskeletal rearrangements that underpin microgamete development.

Electron microscopy was used to bring ultrastructural insight into the **DDD01035881** phenotype. Gametocytes were treated with either compound or DMSO prior to activation and fixed at 25 min post-activation. **Figure 4E** showed an example of a DMSO-treated control gametocytes preparing for exflagellation, forming the characteristic 9+2 organisation of microtubules which emerge as gametes (**Figure 4H**). In contrast, gametocytes treated with **DDD01035881** prior to activation demonstrated a disruption to the structural integrity of the nucleus and food vacuole, with multiple haemozoin-containing vesicles dispersed across the cytosol of the parasite (**Figure 4F**). This finding was consistent with the phenotype of previously described lines with targeted gene disruption of the *Pfs16* gene^30^. The mixed egress phenotype was visualised with gametocytes lacking (**Figure 4F**) and retaining the 4-layer membrane (**Figure 4G**). Again, if **DDD01035881** treatment results in inhibition of microgametogenesis due to Pfs16 binding, the compound could serve as a tool for understanding Pfs16 function, which is yet to be fully elucidated.

Of note, the phenotypes seen with **DDD01035881** treatment by IF were markedly different to inhibition of microgametogenesis when targeting calcium-dependent protein kinase 4 (CDPK4) (using inhibitor 1294^41^), cyclic-GMP dependent protein kinase (PKG) (using the inhibitor ML10^42^) or canonical inhibitors of microtubules (Colchicine) or actin microfilaments (Cytochalasin B) (**Figure S8**). The specific activity windows of 1294 and ML10, 0-20 sec and 0-8 min, respectively, were also markedly different from that of **DDD01035881**, suggesting that compound activity has a discrete function to that of these known regulators of microgametogenesis.

### DDD01035881 treated parasites demonstrate disruption of parasitophorous vacuole and Pfs16 release

Finally, having defined when the compound series is active, we next sought to correlate **DDD01035881** action with the cellular distribution and effects of Pfs16 during microgametogenesis^30^. Pfs16 is known to be localised to the PVM^17,29,38^, which vesiculates and disintegrates prior to the host erythrocyte in an inside-out mechanism of egress during microgametogenesis^37^. Following egress, Pfs16 has been detected in both so-called “Garnham” bodies and multi-laminated whorls formed from the PVM after rupture^38^. By IF labelling of Pfs16, we could show that Pfs16 surrounded microgametocytes at 0 min before either capping or forming a pore at a single side of the activated gametocyte in preparation for egress at around 5.5 min (**Figure 5A**). Upon egress from the host erythrocyte, Pfs16 localised to vesicles which are expelled from the single pore or cap at 6.5 min. Minimal remnants of Pfs16 were seen to remain attached to the parasite from 8.5 min, with no Pfs16 detected on microgametes at 20 min (**Figure 5A**).

**Figure 5.**
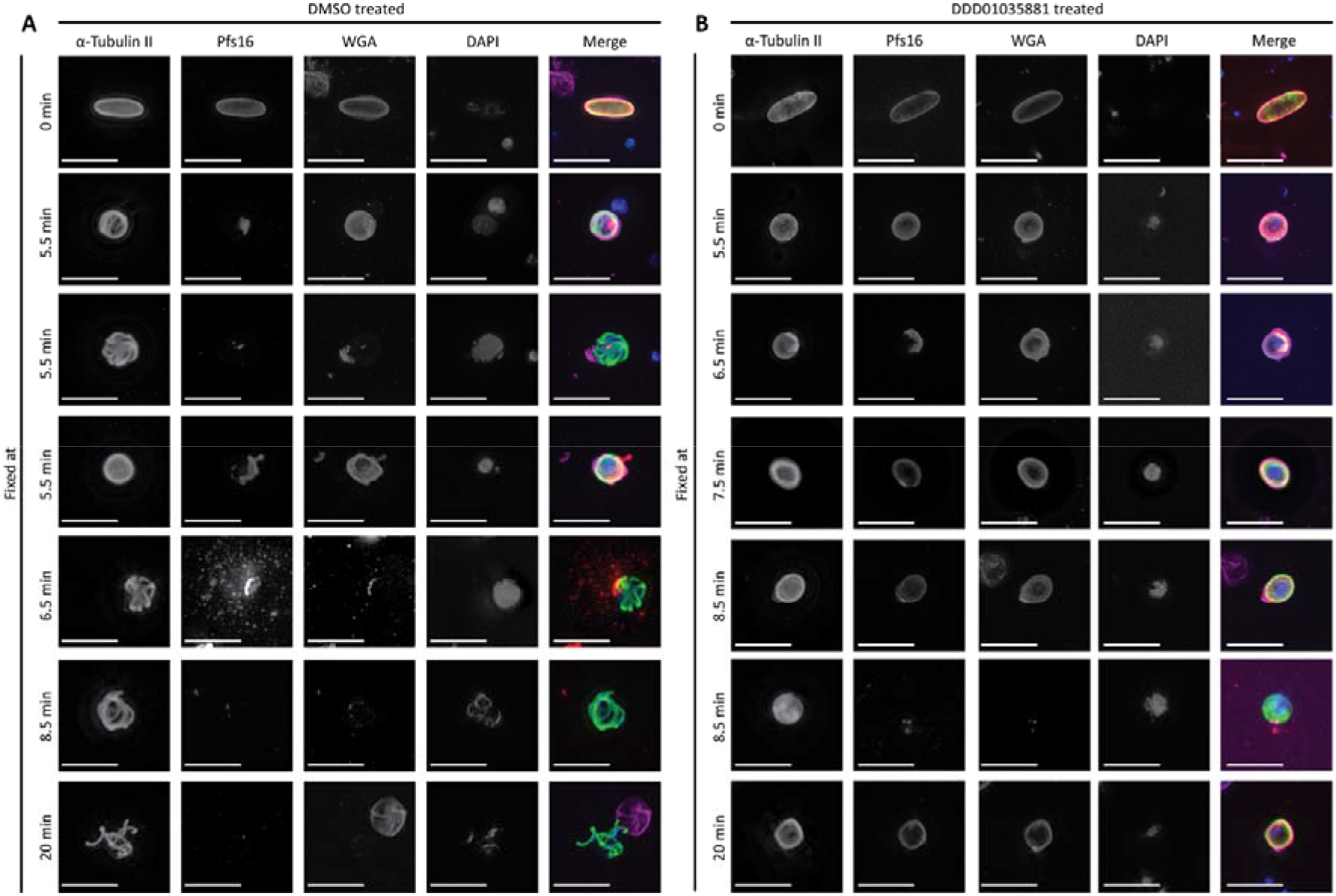
Pfs16 localisation during microgametogenesis with and without DDD01035881 treatment. An IFA time course of microgametogenesis displaying the localisation of PVM protein, Pfs16, as parasites egress from host cells via the inside-out mechanism. Individual channels are displayed on the left of all merged channels, displaying alpha-tubulin (green), erythrocyte membrane (pink), Pfs16 (red) and DNA (blue). Scale bars = 10µm. **(A)** DMSO treated gametocytes. As the PVM prepares to disintegrate prior to erythrocyte egress, Pfs16 either caps or gathers at a pore on the parasite surface at 5.5 minutes post-activation. At the point of egress, the capped or pore-localised Pfs16 vesiculates and bursts at the surface of the parasite at around 6.5 minutes. Beyond the point of PVM and erythrocyte egress, Pfs16 is absent from the parasite. **(B) DDD01035881** treated gametocytes demonstrate a mixed phenotype, with populations of male gametocytes either failing to or successfully egressing from the host cell. Parasites which fail to egress demonstrate continual Pfs16 localisation to the PVM across the entire time course. Parasites which do egress from the host cell demonstrate capping of Pfs16, but Pfs16 does not form a pore, vesicles or burst from the PVM.

At egress, **DDD01035881** treatment was found to disrupt Pfs16 localisation with two distinct phenotypes (**Figure 5B**). In the first, gametocytes failed to egress from the host erythrocyte with Pfs16 retaining expression at the PVM that, as well as the erythrocyte, does not disintegrate or vesiculate. In the second, successful but aberrant erythrocyte egress was detected, as Pfs16 capped to a single end of the parasite but failed to vesiculate and erupt from a pore. With the latter, minimal remnants of Pfs16 were detected at 8.5 min, with no detectable WGA staining. These results demonstrate **DDD01035881** inhibition often disrupts the expulsion of the PVM and associated Pfs16 during microgametogenesis. The combined weight of experimental data therefore not only points to Pfs16 being the target of the N-4HCS scaffold but corroborates Pfs16’s key role in PVM degradation being a critical step in microgametogenesis.

## DISCUSSION

As resistance inevitably threatens the long-term success of artemisinin and its derivatives in treating malaria disease, there is an urgent need for new antimalarials with novel chemotypes and modes of action^43^. Here, we have validated the cellular effects of a potent transmission blocking compound series based on an N-4HCS scaffold and identified the *Plasmodium falciparum* 16 kDa protein, Pfs16, as a highly promising cellular target. The hit compound series was first identified in a cell-based high throughput screen^14^, with hits further optimised by medicinal chemistry^15^. The N-4HCS activity profile fits with the Medicines for Malaria Venture target candidate profile 5 (TCP5) that covers transmission blocking interventions^43^. With identification of its target, and its low cytotoxicity, the N-4HCS scaffold is clearly poised for development as a combination, transmission blocking therapeutic.

Based on detailed phenotypic analysis, the N-4HCS series specifically results in the potent inhibition of microgametogenesis, though without impacting sexual conversion or early gametocyte development. However, the phenotypic effect of the compounds is clear and rapid, traversing the erythrocyte membrane and having action within the first 0-6 min post-activation of parasites. The activity window supports evidence that the Pfs16 protein function is vital to microgametogenesis in its earliest stages, during which the PVM remains associated with the parasite. Beyond PVM expulsion from 6 min, compound effects halt, suggesting that Pfs16 is no longer essential beyond this point. This fits broadly with gene knockout studies for *Pfs16*^30^.

One inconsistency that was noted is that **DDD01035881**-treated parasites did not show any defect in gametocyte commitment, noted as a *Pfs16* knockout phenotype. This may be due to the target site of the Pfs16 protein. A previous study on protein trafficking across the PV to the PVM in *P. falciparum* used parasites transformed with different Pfs16-GFP constructs to determine the amino acid sequences required for PVM targeting and retention^44^. Interestingly, the study suggested the region for PVM targeting and retention differs from the region required for capping during gametogenesis, supporting a dual role for the protein. PVM targeting and retention was shown to be sufficient with inclusion of the 53 C-terminal AA of Pfs16, containing a motif conserved in other known PVM proteins, and N-terminal secretory signal sequence. The specific signal for PVM targeting was localised to 42 AAs comprising the transmembrane domain (22 AA) and part of the C-terminal tail (20 of the 31 AA). The protein membrane interaction was found to be stabilised with 11 C-terminal AA, which when removed reduced the level of retention but did not affect PVM targeting. In contrast, amino acids between the N-terminal secretory signal sequence and transmembrane domain were found to be crucial for capping involved in egress during gametogenesis, though not required for PVM targeting^44^. This difference in function of the different regions of the Pfs16 protein may clarify why the N-4HCS series specifically targets gametogenesis but fails to inhibit commitment to gametocyte formation, despite the protein being present in both stages. Thus, the compound series may potentially bind the region involved in capping; a region suggested to be located within the PV in association with the gametocyte surface. In contrast, the region involved in membrane retention which likely falls on the erythrocyte side of the PVM, a region also shown to be conserved in other PVM proteins, is unlikely to be the target and therefore might explain the specific phenotype of N-4HCS treatment. Further structural investigation of the specific binding of **DDD01035881** to the Pfs16 protein is clearly required to test this hypothesis.

Targeting sexual conversion with antimalarials represents several challenges. Exploiting early commitment or development in a clinical setting is particularly challenging since these parasites develop cryptically in the mammalian host^10^. However, targeting the mature gametocytes to prevent onwards transmission provides several avenues for drug development ^43^. This is where the microgametogenesis targeted-activity of the N-4HCS series may be particularly valuable. The immediate and potent activity of **DDD01035881**, for example, upon activation of gametogenesis (and proven activity in a murine *in vivo* model in this context^14^) gives confidence about the ability of this series to halt parasite development in the mosquito vector. However, for effective inhibition of gametogenesis to occur following a mosquito feed, the compound would need to last long enough in circulation to be taken up by a feeding mosquito. This likely represents the key challenge for this class of molecule. Approaches that may work in this context would be further medicinal chemistry to increase the longevity/bioavailability of this scaffold in the blood stream (current evidence suggests it may only last an hour or so^14^). Alternatively, other approaches to maximise longevity might include formulation for slow release, such as via engineering nanoparticles or other substrates, which facilitate slow release in the blood stream^45^. Recent work has explored impregnation of bed nets or other bait sources with antimalarials to target transmission directing drug uptake to the mosquito not the human host^46,47^. Such an approach may work well with the N-4HCS compound series.

A constant challenge with mode of action identification in *Plasmodium* sexual stages is that these stages of the lifecycle do not replicate. This means that *in vitro* evolution and whole genome analysis (IVIEWGA), a method in which parasites are continually exposed to antimalarial compounds of interest to yield resistance before determining the genetic basis of resistance^48^, will not work^48^. Thus, whilst this chemogenetic approach has been hugely valuable in mapping the druggable genome it remains unattainable for studying targets of compounds which specifically inhibit viability of the sexual stages. This demonstrates the power of CETSA, especially with advancement of protocols specifically for *P. falciparum* drug-target identification^49^. A recent proof-of-concept study used CETSA coupled to mass spectrometry, with both a whole cell and cell lysate approach, to validate targets of pyrimethamine and E64d. Having proven CETSA to be efficacious and robust, the study went on to define to MoA of quinolone drugs, quinine and mefloquine^50^. Although the majority of CETSA studies have been applied to soluble proteins, ligand stabilisation of detergent solubilised membrane proteins has also been successfully achieved^51,52^. An in-depth study on multipass transmembrane proteins utilised live cell CETSA with a range of concentrations of varying detergents, which were added after the compound treatment and thermal challenge of cells^52^. Here, we have demonstrated the power of PAL combined with CETSA in the identification and validation of a PVM cellular target from cellular phenotypic screen-derived hits.

In summary, we have identified Pfs16, described as the earliest marker of sexual conversion, as a very strong candidate for the cellular target of the N-4HCS transmission blocking compound scaffold. Further investment in both the compound but also the protein target itself is now clearly warranted, with the structure of the protein and subsequent attempts at co-crystallisation being a key priority. With further development in the chemistry of transmission blocking drugs and the exploration of avenues to either co-formulate with treatment drugs or deliver via alternative vector targeting approaches, transmission blocking drugs should strongly be considered as important components of future antimalarial combination therapies. With the SARS-CoV-2 pandemic threatening to take resources away from malaria and the looming possibilities of a resurgence of malaria incidence in the developing world, new strategies to break the cycle of infection are more important than ever.

## ACKNOWLEDGEMENTS

We thank Kathrin Witmer, Alisje Churchyard, Irene Garcia-Barbazan, Oriol Llorà-Batlle, Farah Dahalan, Eliana Real and David Grimson, for assisting with parasite culture and for sharing expert transmission blocking advice. Additional thanks to other members of the Baum lab, Fuchter group and Tate lab for assistance with this study. We thank George Ashdown (Baum lab) for assistance in design of imaging experiments, Jane Srivastava and Jessica E Rowley of the Imperial College London flow cytometry facility for assistance in all flow cytometry-based experiments and Ryan Howard and Henry Benns for assisting in the design of photoaffinity labelling experiments. We thank Beatriz Urones Ruano, Ollala Sanz and Sonja Ghidelli-Disse at GSK and Cellzome for helpful discussions and experimental support. We also acknowledge Mathieu Brochet and colleagues for providing assistance in the design of the flow cytometry-based assay for measurements. Finally, we gratefully acknowledge the key role of the Dundee Drug Discovery unit in putting together the original Global Health Chemical Diversity Library (GHCDL) library and generous collaborators who kindly donated parasite lines, additional drugs and antibody reagents critical for completion of this study (Baker, Ishino, Lasonder, Marti, Sauerwein and Van Voorhis labs).

## AUTHOR CONTRIBUTIONS

S.Y., C.N.S, U.S., O.J.F., A.R.Z, S.H., G.V.B., S.J., S.H., M.J.D, E.W.T., A.B., M.J.F and J.B. designed experiments. S.Y., C.N.S, U.S., O.J.F., A.R.Z, S.H., G.V.B. and S.J. performed experiments. S.Y., C.N.S. and J.B. wrote the manuscript. All authors contributed to manuscript preparation.

## DATA AVAILABILITY

Raw proteomics data is attached in the extended supplementary information. Raw files analysed in MaxQuant (version 1.6.1.0) were searched against the curated Uniprot P.falciparum NF54 proteome^53^ using the built-in Andromeda search engine.

The PlasmoDB database (https://plasmodb.org/) was used to analyse protein expression.

## MATERIALS AND METHODS

### In vitro culture of *P. falciparum* NF54 asexual blood stages and gametocytes

*P. falciparum* NF54 strain (sourced from MR4 https://www.beiresources.org/About/MR4.aspx), *PfDyn*GFP/*P47*mCherry (kindly gifted by Edwin Lasonder, Richard Bartfai and colleagues^39^) and *Pf*2004/164-tdTom parasites (kindly gifted by Nicolas Brancucci and Matthias Marti^36^) were cultured for asexual and sexual stage growth as previously described^54^. In brief, asexual blood stage cultures were maintained at 0.75-5% parasitaemia and 4% haematocrit using O+ or A+ human erythrocytes (NHS National Blood Service) supplemented with 30 units/ml heparin (Sigma-Aldrich). Culture medium was prepared from RPMI-1640 with 25mM HEPES (Life Technologies), supplemented with 50 μg/ml hypoxanthine (Sigma), 2 g/l sodium bicarbonate and 10% A+ human serum (Interstate Blood-Bank). CM was changed daily, and cultures were maintained at 37°C under 3% O_2_/5% CO_2_/92% N_2_ gas (BOC Special Gases). For *Pf*2004/164-tdTom parasites, asexual parasites and gametocytes were cultured as described, but maintained at 5% haematocrit in media supplemented with 4nM WR 99210.

Gametocytes were induced from asexual blood stage cultures at 3% parasitaemia and 4% haematocrit. Gametocytes were grown in RPMI-1640 with 25mM HEPES supplemented with 2 mg/ml D-glucose, 150 µg/ml L-glutamine, 2.78 mg/ml sodium bicarbonate, 50 µg/ml hypoxanthine, 5% A+ human serum and 5% AlbuMAX II (Life Technologies). Gametocyte media was changed daily without the addition of fresh erythrocytes for 14 days following induction, at which point stage V gametocytes were most abundant. The functional viability of gametocytes was determined at day 14 post-induction by measuring percentage exflagellation relative to total erythrocyte density. Cultures were activated with ookinete medium (culture medium prepared as above supplemented with 100μM xanthurenic acid, lacking serum or AlbuMAX II) and exflagellation events were counted with a haemocytometer (VWR) using a Nikon Leica DC500 microscope.

### Compounds

**DDD01035881** and **DDD01028076** were purchased from Life Chemicals Inc. and maintained at 10mM in DMSO (Honeywell). **DDD01028076** was also synthesised by ARZ in house^15^. For full methods on the synthesis of clickable derivatives of the N-4HCS series, see **Supplementary Materials and Methods**. ML10 was kindly donated by the Baker lab^42^ whilst BKI-1294 was kindly donated by the Van Voorhis lab^41^. Colchicine and Cytochalasin B were commercially sourced. All compounds were made up to 10mM stock solutions in DMSO (Honeywell) and stored at −20°C.

### Activated gametocyte lysate preparation

Stage V *P. falciparum* NF54 gametocytes were purified by differential sedimentation using NycoPrep™ 1.077 to remove the asexual parasite reservoir. Purified gametocytes were treated with ookinete medium to activate gametogenesis before halting the process at 2 minutes post-activation at 4°C with 0.01% saponin, used to lyse erythrocytes. 5 saponin lysis steps were repeated at 4°C and parasites were washed in PBS before snap freezing in liquid nitrogen and storing at −80°C. Pellets were used in lysate labelling assays and CETSA.

## TARGET IDENTIFICATION

### Lysate Labelling Assays for In-Gel Fluorescence Probe 2 Treatment of Cell Lysate

For in-gel fluorescence detection of probe **2-**treated lysate (**Figure S1A**), 120ml of untreated *P. falciparum* NF54 stage V gametocyte culture was purified using NycoPrep™ 1.077 and saponin lysis as above. The resulting cell pellet was lysed by gentle agitation in 1.2 ml lysis buffer (1% Triton-X100, 10 mM Tris, 150 mM NaCl, 1 x Complete EDTA-free protease inhibitor (Roche Diagnostics)) for 30 min at 4°C. Protein concentration was determined using the BioRad DC Protein Assay, performed according to the manufacturer’s instructions. Absorbance was measured at 750 nm and bovine serum albumin was used as a protein standard. 96-well plates were used to measure absorbance using a SpectraMax M2e Microplate Reader (Molecular Devices). Lysates were made up to 1 mg/mL in lysis buffer, transferred to microcentrifuge tubes and centrifuged (17,000 x *g*, 10 min, 4°C) to remove insoluble cellular debris. Protein was divided across six aliquots of 200 µl before treatment with probe **2** or DMSO, under conditions stated in **Table S2**. Samples were incubated at 4°C for 30 min and irradiated by UV at 254 nm for 5 min.

### Copper catalysed azide-alkyne cycloaddition (CuAAC) and protein precipitation

A click reaction master mix was prepared by combining the following reagents in order:

1. AzTB capture reagent^16^ (1 vol of 10 mM DMSO stock; 0.1 mM final concentration)
2. CuSO_4_ (2 vol of 50 mM H2O stock; 1 mM final concentration)
3. TCEP (2 vol of 50 mM H2O; 1 mM final concentration)
4. TBTA (1 vol of 10 mM DMSO stock; 0.1 mM final concentration).

6 μl of this master mix was added per 100 μl protein sample, vortexed and incubated with moderate shaking for 1 hour at RT. For negative click control samples, H_2_O was added in place of the CuSO_4_ catalyst. The reaction was quenched by addition of 5mM EDTA (from a 500 mM stock in H_2_O). Proteins were precipitated by addition of MeOH (4 vol), chloroform (1 vol) and H_2_O (3 vol) for nLC-MS/MS, or MeOH (2 vol), chloroform (0.5 vol) and H_2_O (1 vol) for gel-based analysis. Precipitated proteins were centrifuged at 17,000 x *g* for 2 min at 4°C. The protein pellet was isolated by removal of the chloroform and MeOH/H_2_O layers and washed with MeOH (4 vol). Protein was then sonicated and transferred to −80°C storage for a minimum of 20 minutes. Samples were centrifuged (17,000 x *g*, 5 min), MeOH was removed and the resulting protein was air dried for 5 min at RT. The protein pellet was resolubilised in 2% SDS and sonicated until fully dissolved before dilution to 0.2% SDS in 1 x PBS solution.

### Pull-down with Dynabeads (Streptavidin MyOne)

Probe **2**-treated and AzTB-labelled cell lysate samples to be analysed by in-gel fluorescence (IGF) were incubated with magnetic Dynabeads (Streptavidin MyOne). Beads were prewashed with 3 vol. of 0.2% SDS, rotary mixing for 3 min at RT. Samples were added to beads and moderately shaken for 2 hours at RT. Supernatant was removed, retaining an aliquot for SDS-PAGE analysis. The beads were then washed with 3 vol 0.2% SDS. Enriched proteins were eluted by boiling with 2% (v/v) 2-mercaptoethanol-containing NuPAGE®LDS sample loading buffer at 95°C for 5 min. Samples were separated by gel electrophoresis.

### Gel electrophoresis and in-gel fluorescence

SDS-PAGE analysis was performed with 12% acrylamide Bis-Tris gels, using a BioRad Mini-PROTEAN® Tetra Cell system with MOPS running buffer (5 mM MOPS pH 7.7, 50 mM Tris Base, 0.1% SDS, 1mM EDTA), using Precision Plus Protein All Blue Standard (BioRad) as a molecular weight marker. IGF was detected (excitation wavelength 552 nm, emission wavelength 570 nm) using a Typhoon FLA 9500 Imager (GE Healthcare). Further data analysis was performed with ImageQuant software.

### Photoaffinity Labelling for nLC-MS/MS

#### Probe 2 Treatment of Live *P. falciparum* Gametocytes

For identification (**Figures 1A-B**, **S1B-C**) and validation (**Figures 1C-F**, **S2**, **S3**) of N-4HCS cellular targets by photoaffinity labelling, live *P. falciparum* NF54 stage V gametocyte cultures (≥ 0.3% exflagellation) were treated and irradiated by UV, as opposed to the lysate-based treatment outlined above. For nLC-MS/MS, gametocytes were treated with either DMSO, probe **2** (10 µM) only or a combination of probe **2** (10 µM) and parent molecule **1** (10 µM). DMSO concentration was normalised across all samples and samples were obtained in triplicate. Following treatment, parasites were incubated for 10 minutes at 37°C and subsequently irradiated with UV light at 254 nm for 10 minutes. Gametocytes were then purified using NycoPrep™ 1.077 and saponin lysed to lyse erythrocytes. Lysis buffer (1% Triton-X100/10mM Tris/150mM NaCl/cOmplete™ ULTRA EDTA-free Protease Inhibitor Cocktail at pH 7.5 in H2O) was added to treated gametocyte pellets and parasites were lysed by sonication (60% amplitude, 3 min, (2 s pulse, 2 s rest)) and centrifugation (17,000 x *g*, 4°C). The supernatant was retained, and probe **2**-labelled proteins were ligated to AzTB by performing the CuAAC reaction as described above.

### Pulldown for nLC-MS/MS

Samples being prepared for nLC-MS/MS analysis were incubated with Neutravidin Agarose beads, which produce a low background signal. For bead derivatisation, beads were centrifuged at 7000 rpm for 4 min at RT before washing 5 times with triethylammonium bicarbonate (TEAB, 100 mM, pH 8). Beads were then gently agitated for 1 hour at RT in a solution of 100 mM TEAB, 25 mM NaBH_3_CN and 0.2% formaldehyde. The reaction was quenched by washing twice with 1% ethanolamine in 100 mM TEAB before subsequently washing 3 times with HEPES (50 mM, pH 8). Before addition of protein samples, derivatised beads were washed twice with 0.2% SDS in HEPES (50 mM, pH 8). Air dried protein samples were dissolved in 0.2% SDS in HEPES (50 mM, pH 8), added to beads and shaken for 2 hours at RT. Following incubation, beads were recovered by spinning at 7000 rpm for 4 minutes at RT and the supernatant was removed. Beads were washed twice with 0.2% SDS in HEPES (50 mM, pH 8) and washed a further 4 times with HEPES (50 mM, pH 8).

### LysC and Trypsin Digestion

To elute protein from beads, LysC (in 50 mM HEPES, pH 8) was added to samples and incubated for 1 hour at 37°C, using 2 µl LysC per 30 µl of derivatised beads. Samples were centrifuged at 6000 rpm 4 min at RT to pellet beads and 50 µl supernatant of samples were retained and combined. Beads were washed with 50 µl HEPES (50 mM, pH 8). TCEP (5 mM) and CAA (10 mM) were added to the combined supernatants and gently agitated 10 min at RT. Trypsin (0.5 µl of 20 µg/100ul in HEPES 50 mM, pH 8.3) was added to each sample and incubated overnight at 37°C.

### 9-plex TMT Labelling

TMT reagents (Thermo Fisher Scientific, MA) were prepared in acetonitrile and added to an equal volume of sample before incubating with moderate shaking for 2 hours at RT. Each reaction was quenched with 1 µl 5% hydroxylamine before combining all samples into one tube. The sample was dried by centrifugal evaporation at 45°C.

### Desalting and 3-Layer Fractionation

All fractionation centrifugations were performed at 1,100 x *g* for 2 min at RT. Samples were resuspended in 150 µl 1% (v/v) TFA/H_2_O, 90% of sample volume was transferred to a stage tip containing SDB-RPS (polystyrene-divinylbenzene copolymer modified with sulfonic acid, Supelco) and centrifuged. Columns were desalted by washing with 0.2% TFA (60 µl) before elution into separate tubes, with sequential addition of three buffers (**Table S3**). Samples were evaporated to dryness in a Savant SPD1010 SpeedVac® Concentrator at 45°C. Prior to separation and analysis by QExactive LC-MS, dried fractionation samples were resuspended in 2% ACN, 0.5% TFA in H_2_O (LC-MS grade) by gental agitation and sonication, to give a final concentration of ∼ 1 µg/µl. A stage tip-filter was prepared, containing 3-layers of PVDF Durapore filter (0.1 µm). Samples (12 µl) were transferred to stage tips and centrifuged into LC-MS vials at 2000 rpm for 3 min at RT.

### nLC-MS/MS Data Acquisition

Peptides were separated on an Acclaim PepMap RSLC column (50 cm x 75 µm inner diameter, Thermo Fisher Scientific) using a 3-hour acetonitrile gradient in 0.1% aqueous formic acid, at a flow rate of 250 nl/min. Easy nLC-1000 was coupled to a QExactive mass spectrometer via an easy-spray source (Thermo Fisher Scientific). The QExactive was operated in data dependent mode with survey scans acquired at a resolution of 70,000 at m/z 200. Scans were acquired from 350 to 1800 m/z. Up to 10 of the most abundant isotope patterns (a minimum of charge 2) from the survey scan were selected with an isolation window of 1.6 m/z and fragmented by HCD with normalised collision energy of 31. The maximum ion injection times for the survey scan and the MS/MS scans (acquired with a resolution of 35,000 at m/z 200) were 20 and 120 ms, respectively. The ion target value for MS was set to 106 and for MS/MS to 2 x 105, and the intensity threshold was set to 1.7 x 103.

### Protein Database Search and TMT-labeling Quantification

Raw files were uploaded into MaxQuant (version 1.6.1.0)^55^ and searched against the curated Uniprot *P.falciparum* NF54 proteome (Uniprot, Feb 2018, 8637 entries)^56^ using the built-in Andromeda search engine. Cysteine carbamidomethylation was selected as a fixed modification, and methionine oxidation and acetylation of protein N terminus as variable modifications. For in silico digests of the reference proteome, the following peptide bond cleavages were allowed: arginine or lysine followed by any amino acid (a general setting referred to as Trypsin/P). Up to two missed cleavages were allowed. The false discovery rate was set to 0.01 for peptides, proteins, and sites. Other parameters were used as pre-set in the software (maximal mass error 4.5 ppm and 20 ppm for precursor and product ions, respectively, minimum peptide length = 7, minimum razor unique peptides = 2, minimum scores for unmodified and modified peptides = 0 and 40, respectively). “Match between runs” option (time window 0.7 min) was allowed and “Unique and razor peptides” mode was selected to allow identification and quantification of proteins in groups (razor peptides are uniquely assigned to protein groups and not to individual proteins), and for TMT quantification (MS2 mode) the minimal ratio count 2 was selected.

### Proteomics Data Analysis

Data analysis was performed using Perseus (version 1.6.5.0)^57^. Corrected reporter intensity values were filtered to remove rows based on ‘contaminants’ and ‘reverse’ columns. The data was log_2_ transformed and the median values within each column (TMT channel) subtracted. Protein groups with at least two valid values were retained. A two-sample t-test (Permutation-based FDR = 0.10; S0 = 0.15) was applied to all proteins in the dataset and results analysed according to their statistical significance.

## TARGET VALIDATION

### Photoaffinity labelling analysed by in-gel fluorescence and western blot

#### Treatment and copper catalysed azide-alkyne cycloaddition

For in-gel fluorescence and western blot analysis (**Figures 1C-F**, **S2**, **S3**), live *P. falciparum* NF54 gametocytes were treated with either DMSO or probe **2** (2.5 µM, 5 µM or 10 µM) and irradiated described above. Gametocytes were purified with Nycoprep 1.077 and saponin lysed to obtain a parasite pellet. The pellets were lysed by sonication (60% amplitude, 3 min, (2 s pulse, 2 s rest)) and centrifuged (17,000 g, 4°C) in lysis buffer (1% Triton-X100/10mM Tris/150mM NaCl/cOmplete™ ULTRA EDTA-free Protease Inhibitor Cocktail at pH 7.5 in H2O).

Protein concentration was determined using the Pierce™ BCA Protein Assay Kit following the manufacturer’s instructions. Absorbance was measured using a NanoDrop™ 2000 spectrophotometer, using BSA as a protein standard. Lysed protein was made up to 0.5-1 mg/ml in PBS to a volume of 100 µl to perform the CuAAC reaction and protein precipitation, as described above. Samples of the clicked sample and crude lysate, with a total of 10 µg protein each, were set aside to be analysed by in-gel fluorescence and western blot.

### Pulldown and IGF

Following CuAAC, protein samples were incubated with Pierce™ streptavidin coated magnetic beads, using 300 µL beads per 1 mg of total protein. Beads were washed 3 times by moderately shaking with 0.2% SDS in PBS and partitioning with a magnet. Protein samples were added to the washed beads and incubated at RT for 2 h, with moderate shaking. The flow-through was retained for analysis by partitioning with a magnet and beads were then washed 3 times with 0.2% SDS. Beads were washed once more with 0.1% Tween-20 in H_2_O before addition of 0.1 M glycine, pH 2.0 and moderate shaking for 5 min at RT. Enriched proteins were then boiled with 2 x sample loading buffer for 10 min at 95°C before centrifuging (17,000 x *g*, 10 min at RT). The resulting eluate was loaded directly onto an SDS-PAGE gel additionally to flow-through, crude lysate and clicked samples.

Proteins were separated on NuPage 4-12% Bis-Tris gels (Novex) and TAMRA IGF was detected with a FLA 5000 biomolecular imager. Gels were further analysed by Western blot using standard methods, developed using ECL (Amersham). Pfs16 was detected with 1:800 mouse anti-Pfs16 clone 32F717:B02^58^, kindly donated by Robert Sauerwein and colleagues, or 1:2000 rabbit anti-Pfg377^59^, kindly donated by Tomoko Ishino and colleagues. Secondary antibodies Goat anti-rabbit or goat anti-mouse horseradish peroxidase (HRP) were used at a 1:10,000 dilution.

### CETSA

The melt curve and isothermal dose response (ITDR)-CETSA protocols were adapted from previously described protocols^34^. An immunoblot-based approach was carried out with activated gametocyte or mixed stage I-III/asexual parasite lysate. The soluble protein fraction of parasites pellets was obtained by addition of lysis buffer (1% Triton-X100/10mM Tris/150mM NaCl/cOmplete™ ULTRA EDTA-free Protease Inhibitor Cocktail at pH 7.5 in H_2_O) and centrifugation at 17,000 x *g* for 20 min at 4°C.

To obtain a melt curve for T_m_ extrapolation, the soluble protein fraction was treated with 1% DMSO or 100µM **DDD01035881**. 10µl aliquots of the protein fractions were aliquoted into PCR tubes and incubated at RT for 3 minutes. The fractions were then thermally challenged over a temperature gradient from either 51.6,76.6, 61.9 or 71.5-100°C for either 5 using the BioRad C1000 Touch™Thermal Cycler and incubated for a further 3 minutes at 4°C. To remove the aggregated proteins, the soluble fraction of the heat-treated parasites was obtained by centrifugation. The stabilised proteins were then prepared for visualisation by immunoblot.

The isothermal dose response format of CETSA was performed over a range of concentrations and single temperature of 78.4°C. The temperature was derived from the melt curve as a temperature at which Pfs16 had mostly aggregated under DMSO treatment but was stabilised by **DDD01035881**. **DD01035881** was dispensed into 384 well plates using the HP D300 Digital Dispenser from 1 nM to 100 µM. 10 µl soluble protein, prepared as described above, was added and incubated with **DDD01035881** for 10 minutes. Treated protein was thermally challenged at 78.4°C for 5-minute and incubated at 4°C for 3 minutes. The stabilised proteins were then isolated from these fractions by centrifugation and prepared for immunoblot analysis.

Gel electrophoresis and detection of Pfs16 and Pfg377 by immunoblot was performed as described above. Densitometry analysis was carried out using ImageJ and data was normalised to the lowest and highest values to obtain relative band intensities. Normalised data was analysed to obtain melt curves using the Boltzmann Sigmoid equation in GraphPad Prism. ITDR data was fitted using the saturation binding curve (one site binding rectangular hyperbola) function in GraphPad Prism.

### Flow cytometry measure of sexual conversion

Sexual conversion rates of *Pf*2004/164-tdTomato parasites was determined as previously described^36^. Asexual parasites with >2% ring-stage parasitemia were synchronised with 5% sorbitol to bring parasites to a window of 0-24 hours post invasion (h.p.i.). Parasites were brought to an 8-hour window (16 to 24 h.p.i) by repeating the synchronisation within the same intraerythrocytic developmental cycle. Synchronised parasites were washed with fresh media, diluted to 2.5% haematocrit and 220µl of dilute cell suspension was plated into the wells of a 96-well plate, for induction of sexual conversion.

25µM of **DDD01035881** and derivatives (**DDD01028076**, **(-)-DDD01028076** and **(+)- DDD01028076**) and DMSO controls were included across the plate, all wells were normalised to 0.25% DMSO. For carryover samples, compounds were plated prior to induction. For post-induction treated samples, compounds were plated 24h after initial plating and induction. For compound washout samples, parasites were plated onto compound treated plates and washed three times at 24 hours-post induction. Parasite controls were included by washing and plating synchronised parasites with media treated with 20µM choline to block sexual conversion.

48 hours after induction, the parasitemia of samples was determined by staining 10µl of each sample with Sybr Green (1:5000 in PBS). Cells were stained for 20 minutes at 37°C and washed twice before measuring Sybr Green positive cells as a measure of parasitemia by flow cytometry using a BD Fortessa flow cytometer. Sybr Green was detected with the 488 nm laser using the 530/30 filter and tdTomato with the 561 nm laser using the 586/16 filter. 100,000 events were acquired per sample. 48 hours after counting parasitemia, Sybr Green staining was repeated to measure gametocytaemia by flow cytometry. Sybr Green and tdTomato double positive cells represented gametocyte populations. 400,000 events were acquired per sample.

The sexual conversion rate of each sample was determined with the following equation:

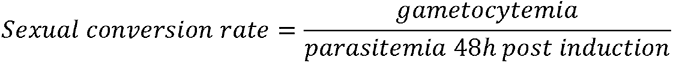

### Relative exflagellation counts

For incremental compound treatment post-gametogenesis, 5 parts of stage V NF54 gametocytes (day 14 post-induction and onwards) were activated with 1-part ookinete medium at RT. Cells were treated with varying concentrations of **DDD01035881**, ML10, 1294 at time increments post-activation. Viability was then determined by counting exflagellation events or cells were prepared for imaging. For viability measurements, exflagellation rates relative to erythrocyte density were determined at 25 minutes post-activation using a haemocytometer. For immunofluorescence labelling, cells were fixed at 25 minutes post-activation and stained as described above.

To determine reversibility of **DDD01035881**, gametocytes were activated as described above and compound was removed with 3 washes at 1 or 6 min post-activation. Viability was determined by counting exflagellation rates at 25-minutes post-activation. Exflagellation rates of **DDD01035881**-treated gametocytes was determined relative to DMSO controls. Relative rates of **DDD01035881**-treated parasites which were washed at 1 or 6 minutes were calculated relative to DMSO controls which were washed three times at 1-minute post-activation, accounting for reductions in exflagellation due to centrifugation.

### Flow cytometry measure of microgametocyte ploidy

The extent of DNA replication of *PfDyn*GFP/*P47*mCherry gametocytes with and without **DDD01035881** treatment was determined as previously described^60^. Stage V *PfDyn*GFP/*P47*mCherry and NF54 gametocytes were purified with NycoPrep™ 1.077. Purified gametocytes were resuspended in suspended activation medium (RPMI-1640 with 25mM HEPES (Life Technologies), 4mM sodium bicarbonate, 5% fetal bovine serum, pH 7.20) to permit staining at RT without premature activation of gametogenesis. For T=15 min samples, parasites were activated with ookinete medium and gametogenesis was halted with ice cold PBS at 15 minutes. For T=0 min samples, parasites were immediately resuspended in ice cold PBS. All samples were washed at 300*g* for 2 min at 4°C, resuspended in ice cold PBS and stained with 1:2000 Vybrant™ DyeCycle™ Violet (Thermo-Fisher) for 30 min at 4°C. Stained and unstained erythrocyte and NF54 (T=0 and T=15 min) controls were prepared. DNA content was measured as Vybrant™ DyeCycle™ Violet intensity by flow cytometry and data was analysed using FlowJo software. GFP positive male gametocytes were gated and ploidy was measured and expressed as a percentage of the total male population. The 530/30, 610/10 and 450/50 filters were used to analyse GFP, mCherry and Vybrant DyeCycle™ Violet, respectively. 100,000 events were analysed per sample.

### Immunofluorescence staining and imaging

Mature NF54 gametocyte culture was treated with either DMSO or test compounds. Gametocytes were treated with 5µM **DDD01035881**, 1µM 1294 and 25µM ML10, before immediately activating without prior incubation. Gametocytes were treated with 50µM Colchicine and Cytochalasin B for 48 hours. For T=0 samples, compounds were immediately fixed in prewarmed 4% paraformaldehyde. Cultures were activated by xanthurenic acid-containing ookinete medium and fixed at increments post-activation. All fixed samples were adhered to poly-L-lysine (Sigma) coated glass coverslips, before cells were washed once in PBS, permeabilised in 0.1% Triton-X100, washed thrice more in PBS and blocked with 10% fetal bovine serum. Cells were labelled with primary antibodies for 1 hour; 1:500 mouse anti-alpha tubulin clone DM1A (Sigma), 1:1000 rabbit anti-Glycophorin A clone EPR8200 (Abcam) and 1:800 mouse anti-Pfs16 clone 32F717:B02^58^ (a kind gift from Robert Sauerwein, Radboud University Medical Centre). Cells were labelled with secondary antibodies and other stains for 45 minutes; 1:500 anti-mouse or anti-rabbit Alexa Fluor 488 (Thermo-Fisher), anti-mouse or anti-rabbit Alexa Fluor 594 (Thermo-Fisher), 5µg/ml Wheat Germ Agglutinin (WGA)-633 and 10nM 4’,6-diamidino-2-phenylindole (DAPI). VectaShield mountant (Vector Laboratories) was used to mount coverslips onto glass slides. Images were acquired with a Nikon Ti-E inverted widefield microscope at x100 objective in 0.2 µm increments through Z, using NIS Elements v4.20. Z-stack images were deconvolved using the EpiDemic plugin and compressed to maximum intensity projections in Icy Bioimage Analysis software.

### Electron Microscopy

Stage V NF54 gametocytes were purified by density barrier isolation with NycoPrepc 1.077. Purified gametocytes were treated with either DMSO or 5µM **DDD01035881** prior to activation with ookinete medium. Parasites were fixed at 25 min post-activation with 4% high EM grade PFA/2.5% v/v glutaraldehyde/ 0.1% tannic acid in 0.1 M sodium cacodylate buffer pH 7.2 for 3 hrs at RT and washed three times in cold 0.1 M sodium cacodylate buffer at 20 min intervals. Cells were treated with 1% w/v osmium tetroxide in 0.1 M sodium cacodylate for 2 hrs at room temperature (RT), washed with 0.1 M sodium cacodylate and stained with 1% w/v aqueous uranyl acetate for 1 hr at RT. Samples were then dehydrated in an ethanol series and embedded in epoxy resin (TAAB). 70 nm sections were cut using a Leica EM UC7 ultramicrotome, contrasted with Uranyless (TAAB) for 2 min and with 3% Reynolds lead citrate (TAAB) for 1 min according to the manufacturer’s protocols. Sections were imaged on a JEOL JEM-1400Plus TEM (120 kV) with a Ruby Camera (2 K × 2 K).

