## Supplementary Information for "*Plasmodium falciparum* protein Pfs16 is a target for transmission-blocking antimalarial drug development"

- **Supplementary Tables**
- **Supplementary Figures**
- **Supplementary Methods**
- **Supplementary References**

#### SUPPLEMENTARY TABLES

| Structure | Compound | Male $IC_{50}$ (nM) |
|---|---|---|
| 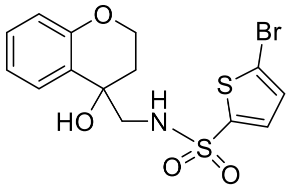 | **DDD01035881** | 292±76 |
| 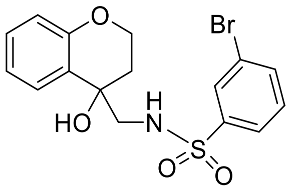 | **DDD01028076**<br>**(-)-DDD01028076**<br>**(+)-DDD01028076** | 367±60<br>18±3<br>2861±325 |
| 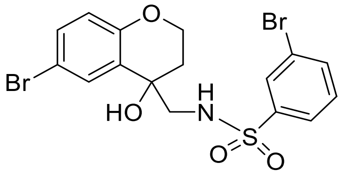 | **1** | 785±48 |
| 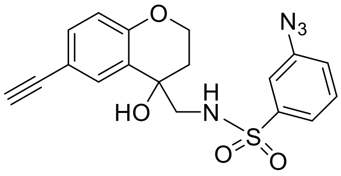 | **2** | 3981±211 |
| 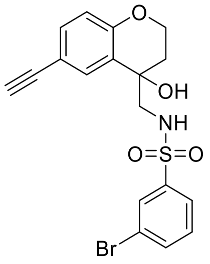 | **3** | 370±79 |
| 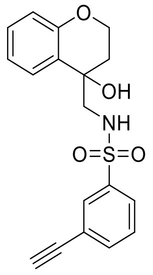 | **4** | >1000 |
| 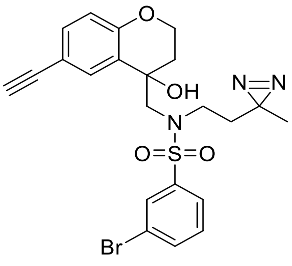 | **5** | >25,000 |

| | | |
|---|---|---|
| 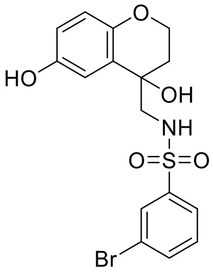 | **6** | >11,000 |
| 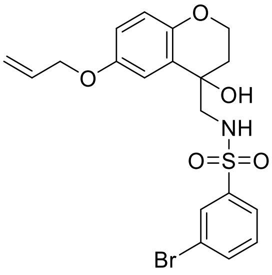 | **7** | >25,000 |
| 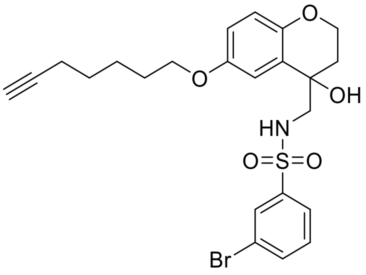 | **8** | >1000 |
| 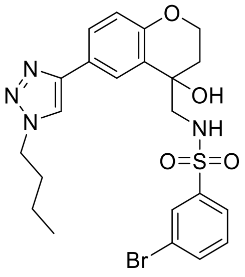 | **9** | >25,000 |
| 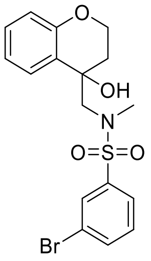 | **10** | >10,000 |
| 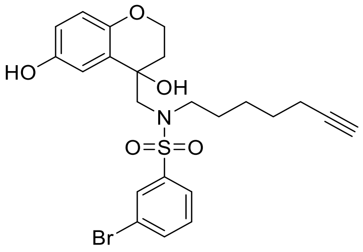 | **11** | >25,000 |

| | | |
|---|---|---|
| 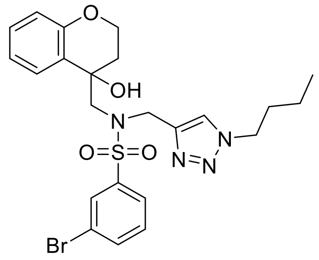 | **12** | >25,000 |
| 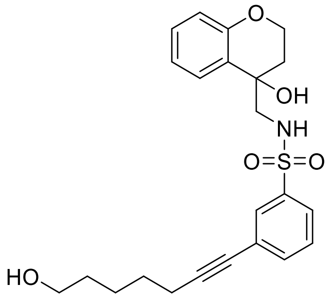 | **13** | >25,000 |

**Table S1. Activity of DD01035881 derivatives towards clickable analogues**

Activity of **DDD01035881** and structural analogues, synthesised to determine functional groups amenable to change and addition of clickable moieties (aryl azide or diazirine). $IC_{50}$s depicted represent the inhibition of male gamete viability as determined by the male gamete formation format of the DGFA. Values are expressed as an average ± SEM of ≥3 biological replicates, ≥ 2 technical replicates [1].

| | Probe | Volume added (uL) | AzTB |
|---|---|---|---|
| A | DMSO control | 10 | + |
| B | Probe **2** 10 µM | 10 | - |
| C | Probe **2** 1 µM | 1 | + |
| D | Probe **2** 2 µM | 2 | + |
| E | Probe **2** 5 µM | 5 | + |
| F | Probe **2** 10 µM | 10 | + |

**Table S2. Probe volumes added to lysate aliquots.**

| | |
|---|---|
| SDB-RPS (1) | Ammonium Formate (100 mM), 40% ACN (v/v), 0.5% Formic Acid (v/v) |
| SDB-RPS (2) | Ammonium Formate (150 mM), 60% ACN (v/v), 0.5% Formic Acid (v/v) |
| Buffer | 5% Ammonium Hydroxide (v/v), 80% ACN (v/v) |

**Table S3. Fractionation elution buffer table.**

#### SUPPLEMENTARY FIGURES

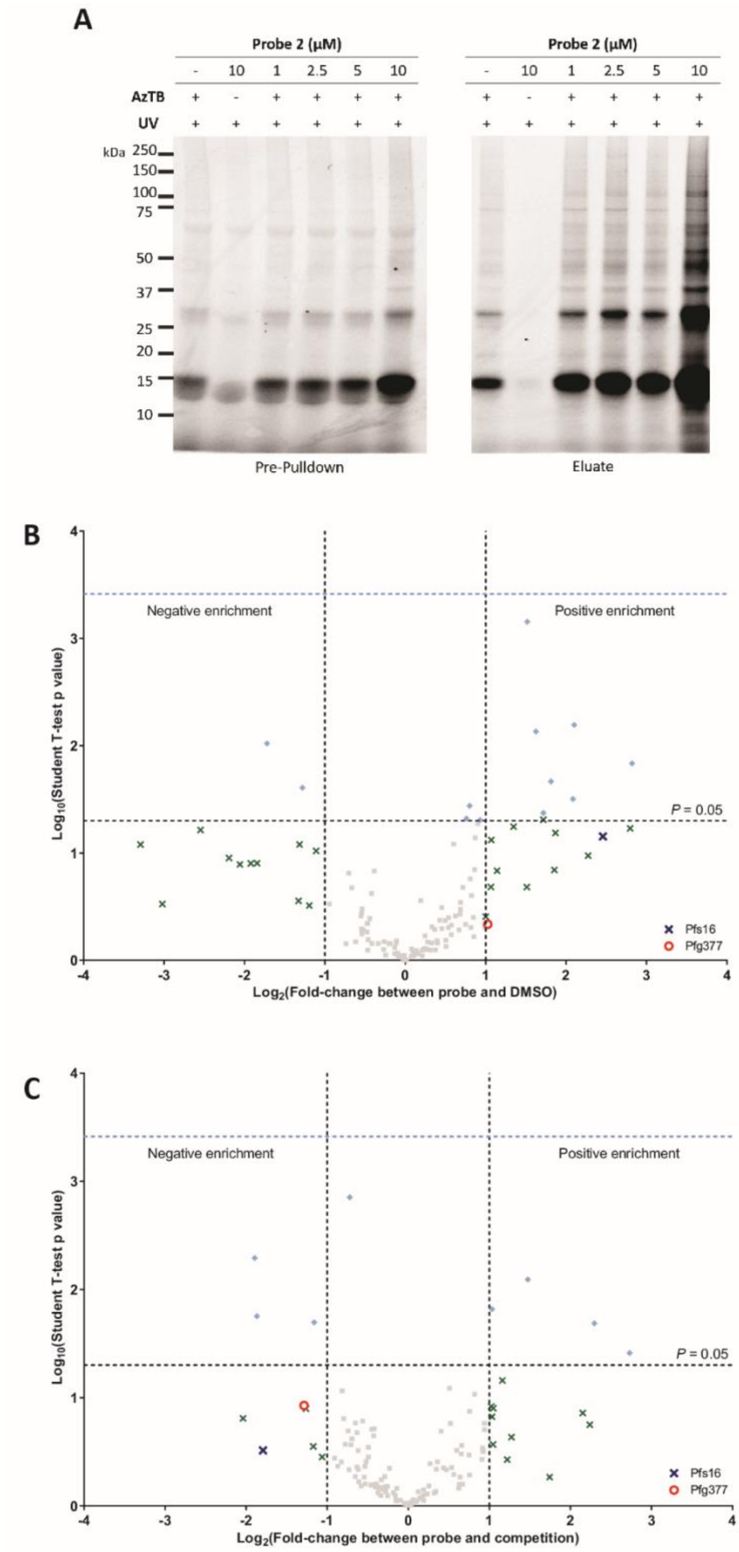


**Figure S1. (A)** Confirmation of protein pulldown in the presence of AzTB capture reagent and probe **2** using cell lysate derived from *P. falciparum* gametocyte culture. Results confirm an AzTB-specific and probe **2** concentration-dependent pulldown by IGF. **(B-C)** Proteome-wide results from live treatment of *P. falciparum* stage V gametocyte analysed in a 9plex TMT study. Plots depict

$log_{10}$-transformed student's t-test p values against $log_2$-transformed fold change in peptide-hit enrichment between **(B)** probe **2** and DMSO treated samples and **(C)** probe **2** and competition (combined treatment probe **2** and parent molecule **1**) treated samples. Fold change was calculated between the average enrichment of 3 biological replicates, per condition. The p value of 0.05 is marked by the horizontal grey dashed line. Green crosses represent proteins with a log-transformed difference in enrichment either above 1 or below -1, for positive and negatively enriched proteins, respectively. Enriched proteins with a p value $< 0.05$ are depicted as light blue diamonds. Grey squares represent the proteins not fitting the enrichment and *p* value thresholds. Pfs16 is marked with a blue cross and Pfg377 is marked with a red circle.

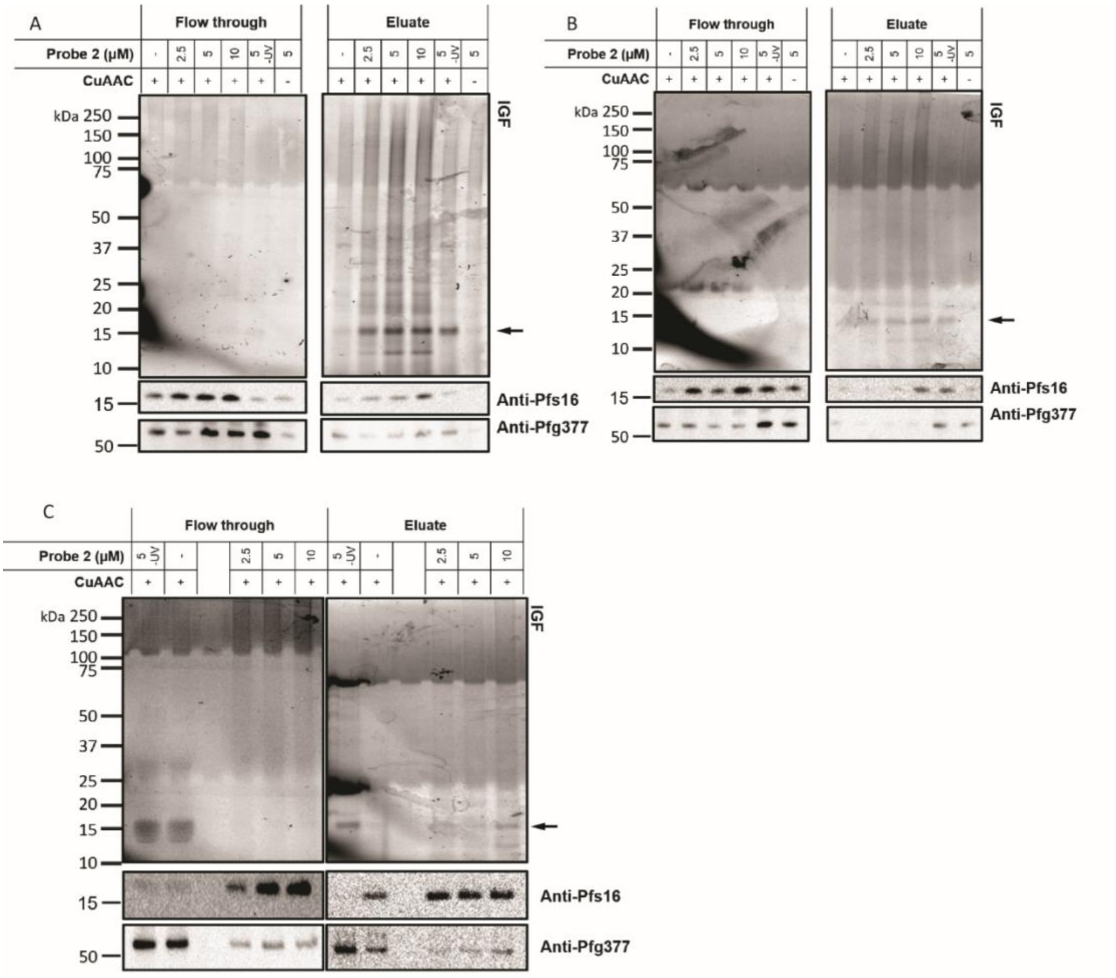


**Figure S2. Live parasite PAL full IGF and corresponding immunoblot data**

**(A-C)** Replicates of raw data acquired from treatment of live gametocytes with increasing probe **2** doses prior to cell lysis, ligation to AzTB and streptavidin-enrichment of labelled proteins. The full IGF and corresponding immunoblots against Pfs16 and Pfg377 depict the flow through and enriched eluate samples from pulldowns.15-20kDa bands of the IGF gels, likely corresponding to Pfs16, and Pfs16 and Pfg377 labelled in immunoblots were analysed by densitometry. CuAAC denotes performance of the click reaction, samples denoted with a – represent click controls lacking a copper catalyst. -UV samples represent UV controls which were not irradiated with UV after probe **2** treatment.

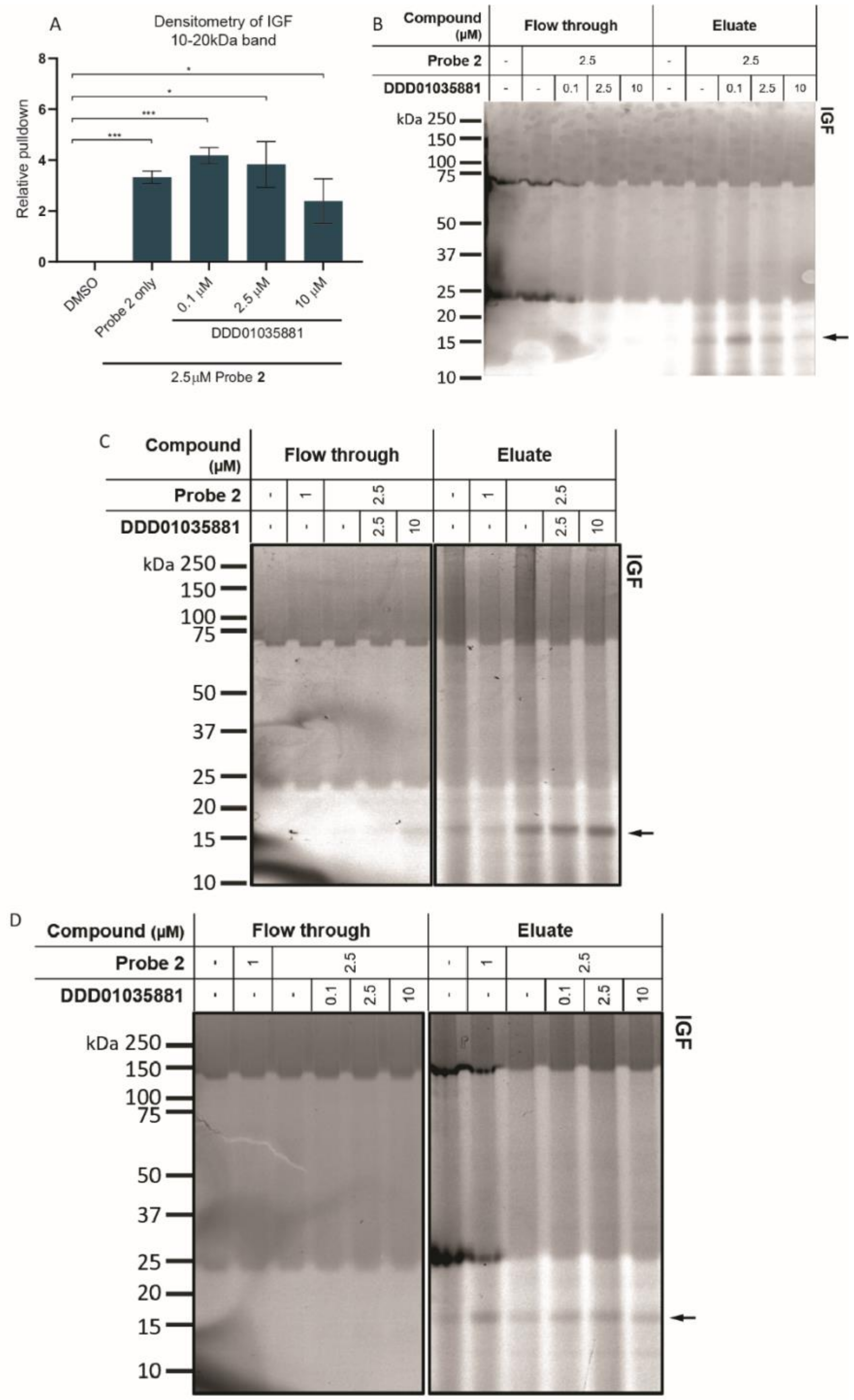


**Figure S3. IGF and corresponding densitometry of live parasite PAL performed in competition with DDD01035881**

Target validation by live treatment with probe **2** and competitor, AzTB conjugation and streptavidin-biotin affinity enrichment. Increasing concentrations of competitor were incubated with a single concentration of probe **2**. **(A)** Densitometry of the 15-20kDa protein band, likely corresponding to Pfs16, in the streptavidin enriched fractions depicted as relative band intensity, relative to a DMSO control. Error bars denote SEM of 2-3 biological replicates. Significance in unpaired two-tailed t-test denoted as * ($p < 0.05$), ** ($p < 0.01$) and *** ($p < 0.001$). **(B-D)** Replicates of raw IGF data analysed for **A** with parent compounds **DDD01035881** (denoted as DDD_881 in **C** and **D**). The 16kDa protein band analysed by densitometry is labelled.

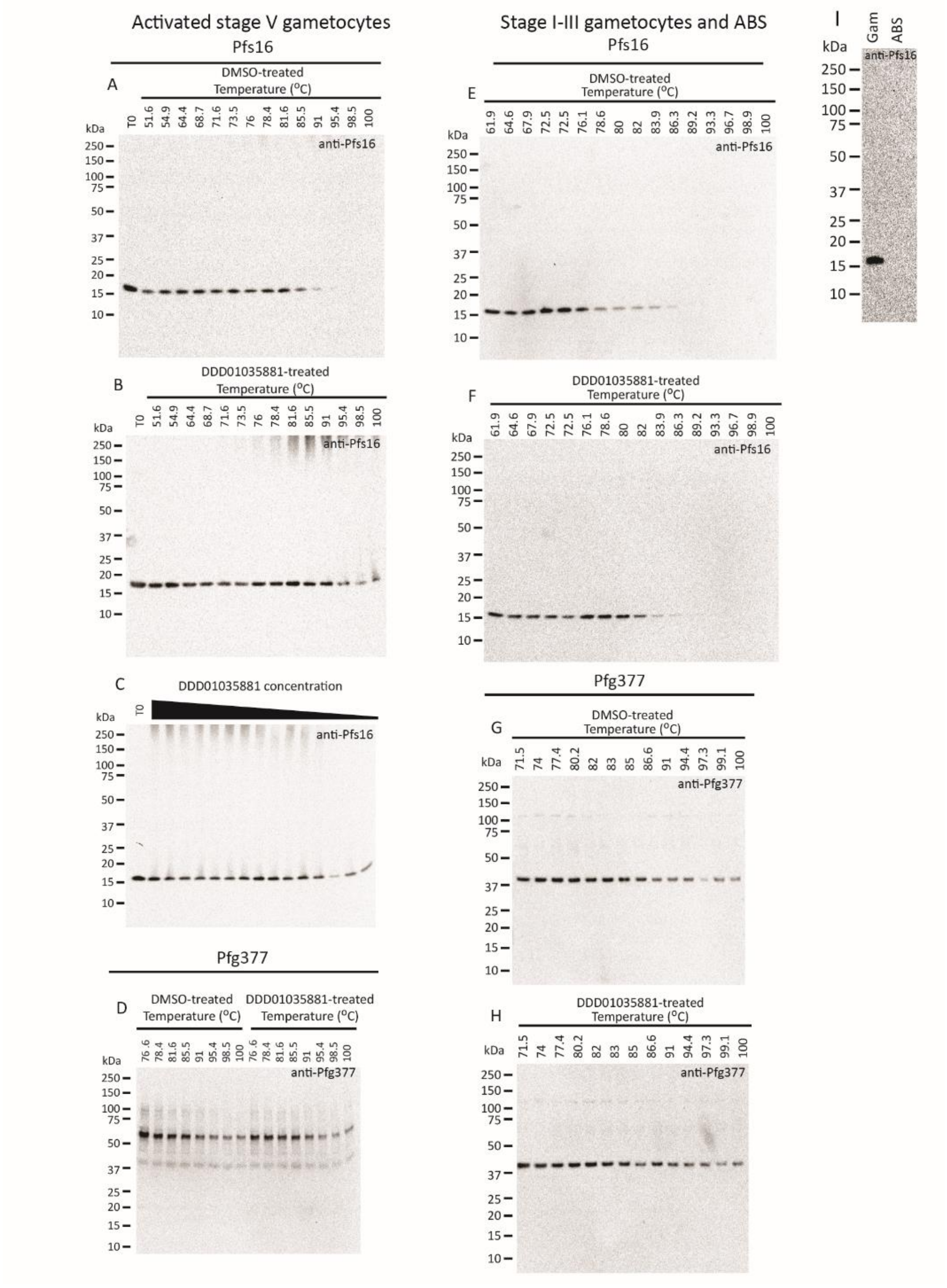


**Figure S4. Immunoblotting for CETSA melt curve analysis**

Full immunoblots obtained for densitometric analysis using activated stage V gametocyte lysate, treated with **(A)** DMSO or **(B) DDD01035881**, thermally challenged from 51.6-100°C and labelled with anti-Pfs16 antibody. **(C)** Anti-Pfs16 labelled immunoblot depicting the ITDR **DDD01035881**-treatment of activated stage V gametocyte lysate from 100µM-1nM and thermally challenged at 78.4°C. T0 controls represent stage V gametocyte lysate. **(D)** Full immunoblot of activated stage V gametocyte lysate protein control labelled with anti-Pfg377, treated with DMSO or **DDD01035881** and thermally challenged from 76.6-100°C. Full immunoblots obtained for densitometric analysis using mixed ABS and stage I-III gametocyte lysate, treated with **(E)** DMSO or **(F) DDD01035881**, thermally challenged from 61.9-100°C and labelled with anti-Pfs16 antibody. Full blot of protein control labelled with anti-Pfg377, treated with **(G)** DMSO or **(H) DDD01035881** and thermally challenged from 71.5-100°C. Note, probing with anti-Pfg377 in activated stage V gametocytes contains an additional uncharacterised band by immunoblot, present at 60 kDa, which is only faintly recognised in ABS or stage I-III gametocytes. **(I)** Full immunoblot demonstrating the gametocyte specific Pfs16 expression, permitting the production of mixed ABS and stage I-III gametocyte lysate used in **E-H** without gametocyte purification.

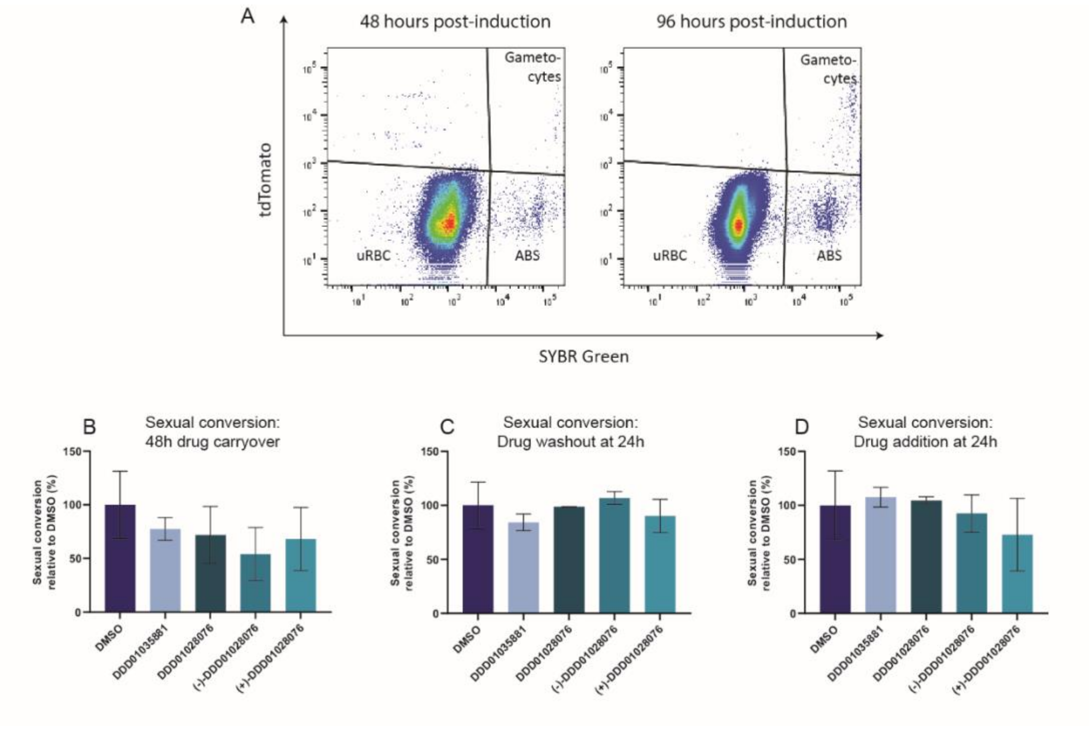


**Figure S5. Flow cytometry of SYBR Green stained *Pf*2004/164-tdTomato gating strategy and N-4HCS analogue results**

**(A)** Sexual conversion rates of *Pf*2004/164-tdTomato parasites were quantified as the tdTomato and SYBR Green positive gametocytes at 96 hours post-induction relative to the parasitemia at 48 hours post-induction, measured as the SYBR Green positive parasite population. Perturbation effects were then quantified relative to DMSO controls. **(B-D)** Corresponding sexual conversion rates of Pf2004/164-tdTomato parasites of **DDD01035881**, **DDD01028076**, **(-)-DDD01028076** and **(+)-DDD01028076**-treated parasites are expressed as rates relative to the conversion rates of DMSO-treated parasites. Perturbations to sexual conversion were determined by **(B)** maintaining treatment over two intraerythrocytic cycles and **(C)** the reversibility of any perturbations were determined by removing compound at 24 hours. **(D)** Perturbations to early gametocyte development were probed by administration of compounds in a subsequent intraerythrocytic cycle. Error bars represent the SEM of 2 biological replicates.

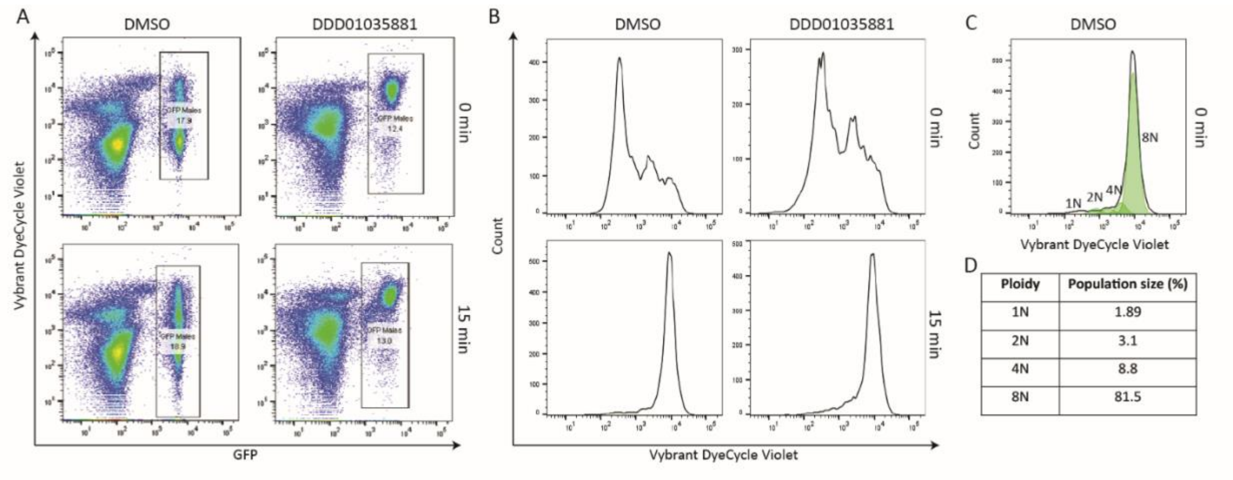


**Figure S6. Flow cytometry of Vybrant™ DyeCycle™ Violet stained *PfDyn*GFP/*Pf47*mCherry gating strategy**

Perturbations to ploidy during gametogenesis were quantified by flow cytometry by **(A)** gating GFP-positive male gametocytes from the 2 conditions of DMSO or **DDD01035881** treatment at 0 and 15-minutes post-activation. **(B)** Histograms of Vybrant™ DyeCycle™ Violet staining of males gated from **(A)** were further gated into 1N, 2N, 4N and 8N populations using the **(C)** proliferation model in FlowJo, before dividing into **(D)** relative populations.

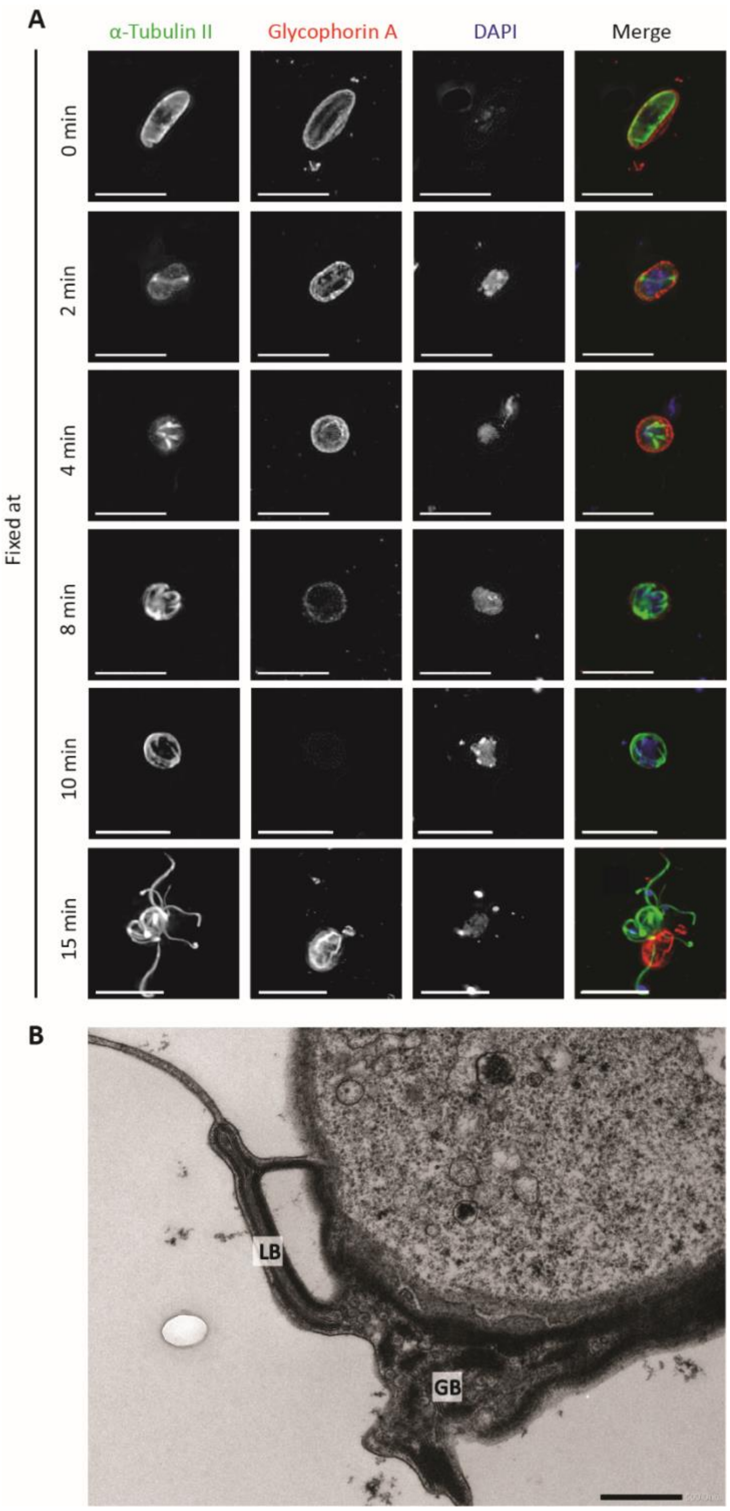


**Figure S7. *P. falciparum* microgametogenesis**

Characterisation of *P. falciparum* (NF54) microgametogenesis *in vitro* which marks the immediate life cycle progression that occurs in the mosquito midgut following transmission from host to vector. **(A)** An IFA time-course of microgametogenesis, depicting individual channels and the merge of alpha tubulin-  labelled cytoskeleton (green), glycophorin A-labelled host erythrocyte (red) and

DNA (blue). Drastic cytoskeletal rearrangement occurs across the entirety of the process as parasites egress from the host erythrocyte and replicate their genome three times, alternating with three endomitotic divisions. Scale bars = 10µm. **(B)** Electron microscopy of activated male gametocyte fixed at 25 minutes post activation. Distinctive cellular features of intraerythrocytic *P. falciparum* gametocytes, the Laveran's bib (LB) and Garnham bodies (G).

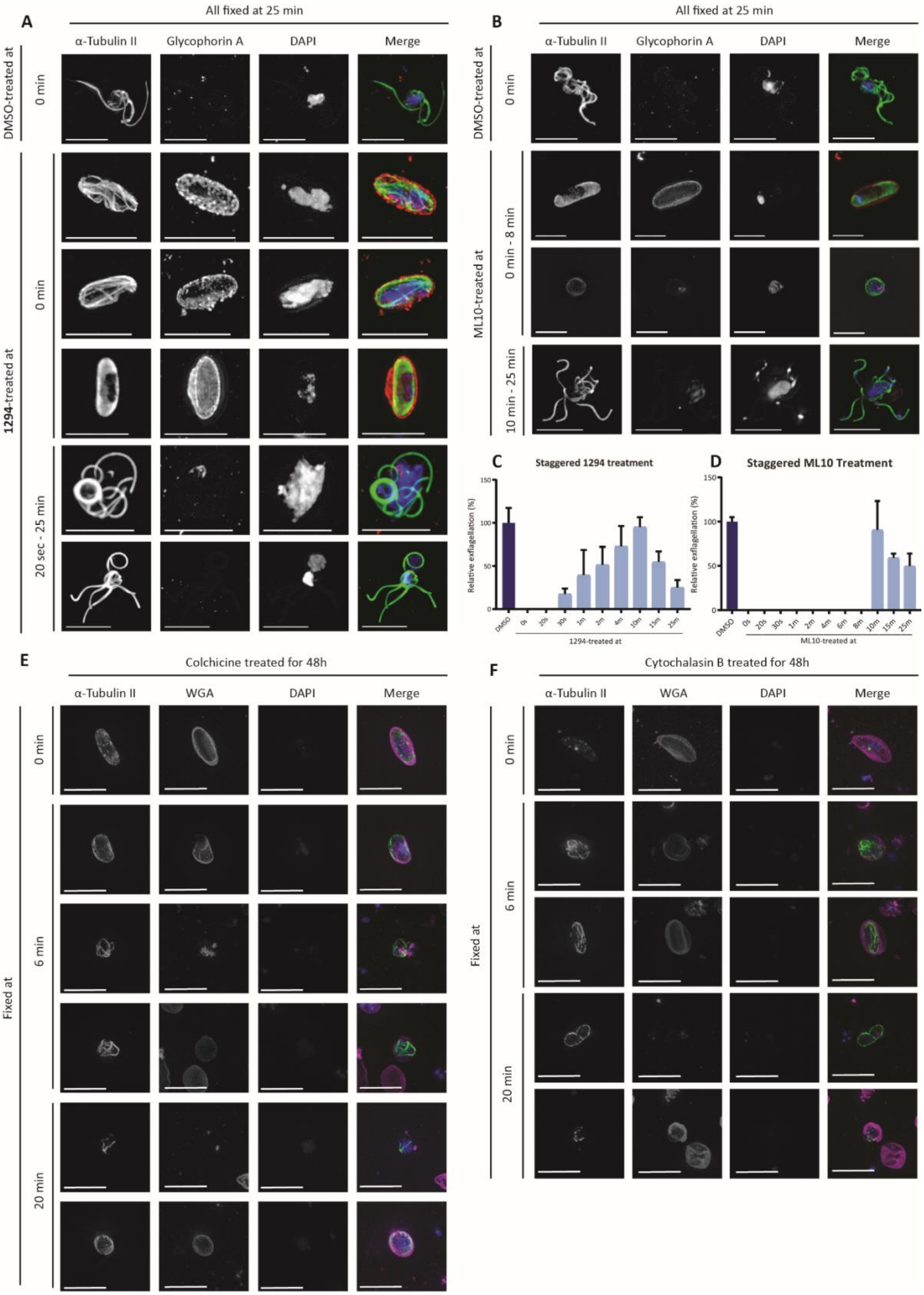


**Figure S8. Phenotype of known small molecular inhibitors of *P. falciparum* microgametogenesis**

IFAs depicting phenotypes of **(A)** 1294, **(B)** ML10, **(D)** Colchicine and **(E)** Cytochalasin B during microgametogenesis. Individual channels of alpha tubulin-labelled cytoskeleton (green), glycophorin A-labelled (red) or WGA-labelled (far-red) host erythrocyte and DNA (blue) are shown alongside merged channels. Scale bars = 10µm. Parasites treated with **(A)** 1294 and **(B)** ML10 were activated, treated in time increments and fixed at 25 minutes, the point DMSO controls exflagellate. **(A)** 1294-treated gametocytes showed one of two phenotypes within the active window; i) gametocytes failed to develop at all, failing to replicate DNA, remaining falciform and intraerythrocytic, or ii) gametocytes failed to round up or egress, but host erythrocytes appeared fragmented, gametocytes replicated DNA and formed elongated axonemes which coiled around the falciform cell body. **(B)** ML10-treatment resulted in two distinct phenotypes; 1) gametocytes demonstrated no morphological progression and retained the morphology of intraerythrocytic stage V gametocytes which failed to fully replicate DNA or exflagellate or 2) gametocytes failed to exflagellate but rounded up and egressed from the host erythrocyte. Phenotypes are grouped according to the distinct phenotypes observed across 30 second-2-minute increments within the stated time windows. Exflagellation rates of gametocytes treated with **(C)** 1294 or **(D)** ML10 relative to DMSO controls when activated and treated at the stated time points. Error bars represent the SEM of 3 biological replicates. Gametocytes were treated with **(E)** Colchicine and **(F)** Cytochalasin B for 48 hours before activating and fixing at the stated timepoints relative to activation. **(E)** Colchicine treatment resulted in impaired axoneme formation and organisation but did not affect DNA replication. **(F)** Cells treated with Cytochalasin B exhibited impaired axoneme assembly and either rounded up without DNA replication or remained falciform but replicated DNA.

### SUPPLEMENTARY METHODS

#### GENERAL SYNTHESIS METHODS

All chemicals were purchased from Sigma-Aldrich Ltd, Fluorochem Ltd., Acros Organics and used without further purification. All reactions were performed under nitrogen or argon atmosphere using dried glassware. Silica gel column flash chromatography was performed using high-purity grade silica gel, pore size 60 Å, 220-440 mesh particle size, 35-75 µm particle size.

Analytical and preparatory HPLC runs were performed on a Agilent 1200 series system with a G1315D detector, a G1361A preparative pump, a semi-prep Daicel Chiralpak IE column 10×250 mm, 5 µm and eluting with 60:40 hexanes:EtOAc, 10 mL/min. For prep scale, 5–10 mg was injected at a time.

NMR spectra were recorded on 400 MHz Bruker instruments at room temperature and were referenced to residual solvent signals. Data are presented as follows: chemical shift, multiplicity (br s = broad singlet, s = singlet, d = doublet, t = triplet, q = quartet, m = multiplet) and integration.

The purity of compounds was verified by $^1$H-NMR as well as RP-HPLC on a Waters 2767 system equipped with a photodiode array and an ES mass spectrometer. Separation was achieved using a XBridge C18 (5 µm, 4.6 mm × 100 mm) column, equipped with an XBridge C18 guard column (5 µm, 4.6 mm × 20 mm) eluting with a gradient of $H_2O$/ACN. Purity of tested compounds was ≥ 95%, unless specified.

Mass spectrometry was performed using chemical ionisation (CI), electron ionisation (EI) or electrospray ionisation on an AUTOSPEC P673 spectrometer by the Chemistry Department Mass Spectrometry Service at Imperial College London.

Unless otherwise stated all reactions were performed using anhydrous solvents, reacted under an atmosphere of argon, monitored by TLC and stirred at RT until completion.

#### SYNTHESIS OF PARENT MOLECULE 1

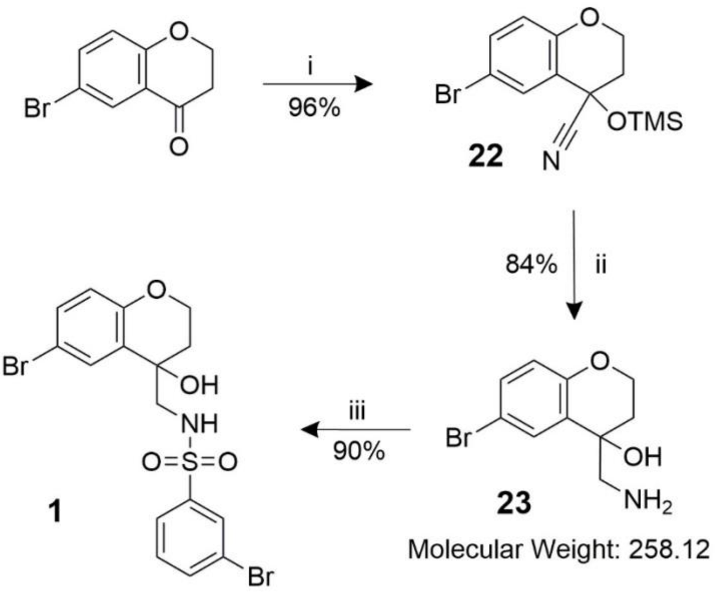


**SCHEME 1: Synthetic route to affinity pulldown inhibitor, Compound 1. i** - $ZnI_2$, TMS-CN, 50°C, ON. **ii** - $LiAlH_4$, THF, 0°C, 3 h. **iii** - TEA, DCM, RT, 5 h.

**Compound 22** - 6-bromo-4-((trimethylsilyl)oxy)chromane-4-carbonitrile

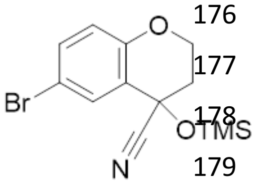


A solution of 6-bromochroman-4-one (454 mg, 2 mmol) and $ZnI_2$ (32 mg, 0.1 mmol) were suspended in DCM and cooled to 0 °C. Trimethylsilyl cyanide (0.38 mL, 3 mmol) was added dropwise and the resulting solution was stirred overnight. The resulting solution was diluted with DCM (20 mL), washed with $NaHCO_3$ (3 × 20 mL), extracted with DCM (2 × 60 mL), dried over $MgSO_4$ and concentrated *in vacuo* to afford the title compound, as an orange oil (626 mg, 96%). Due to column instability, compound **22** was carried forward without further purification

$^{1}$H NMR (400 MHz, Methanol-$d_4$) δ 0.23 (s, 9H), 2.40 (ddd, J = 13.9, 5.5, 3.4 Hz, 1H), 2.49 (ddd, J = 14.0, 8.7, 4.4 Hz, 1H), 4.31 – 4.43 (m, 2H), 6.83 (d, J = 8.9 Hz, 1H), 7.45 (dd, J = 8.8, 2.5 Hz, 1H), 7.63 (d, J = 2.5 Hz, 1H). $^{13}$C NMR (101 MHz, Methanol-$d_4$) δ -0.23, 35.47, 61.51, 65.34, 111.88, 119.49, 120.32, 123.07, 130.51, 133.91, 152.88. TLC (hexane: EtOAc – 3:1) Rf: 0.59

**Compound 23** – 4-(aminomethyl)-6-bromochroman-4-ol

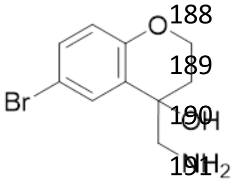


A suspension of $LiAlH_4$ (303 mg, 8 mmol) in THF was cooled to 0°C. Compound **22** (652 mg, 2 mmol) was dissolved in THF and added drop-wise over 15 min. The reaction was stirred at 0°C until completion. The reaction was quenched using Fieser procedure, the following solutions were added slowly at 0°C: dilute with $Et_2O$, $H_2O$ (1 × w/v mass of $LiAlH_4$), 15% NaOH (1 × w/v mass of $LiAlH_4$), $H_2O$ (1.5 × w/v mass of $LiAlH_4$). The solution was warmed to RT, stirred for 15 min, dried over $MgSO_4$, filtered and concentrated *in vacuo*. This afforded the tile compounds as a clear oil (433 mg, 84%)

$^1$H NMR (400 MHz, Methanol-$d_4$) δ 1.98 (ddd, J = 14.0, 8.1, 4.1 Hz, 1H), 2.23 (ddd, J = 14.1, 6.7, 3.6 Hz, 1H), 2.93 (q, 2H), 4.18 – 4.30 (m, 3H), 6.73 (d, J = 8.8 Hz, 1H), 7.27 (dd, J = 8.8, 2.5 Hz, 1H), 7.57 (d, J = 2.5 Hz, 1H). $^{13}$C NMR (101 MHz, Methanol-$d_4$) δ 31.67, 50.16, 63.24, 67.92, 111.96, 118.52, 129.31, 129.36, 131.39, 153.89. MS: *m/z* (ES) 240 (50%, $[M-H]^+$), 258 (100%, $[M+H]^+$). HRMS, found 258.0134 ($C_{10}H_{13}NO_2Br$, $[M+H]^+$, requires 258.0130). TLC (DCM:MeOH – 90:10) $R_f$: 0.19.

**Compound 1** – 3-bromo-N-((6-bromo-4-hydroxychroman-4-yl)methyl)benzenesulfonamide

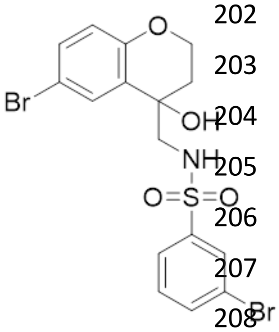


Compound **23** (26 mg, 0.10 mmol) was dissolved in DCM under an inert atmosphere at 0 °C, before Triethylamine (TEA) (0.04 mL, 0.25 mmol) was added dropwise. A solution of 3-bromobenzenesulfonyl chloride (33 mg, 0.13 mmol) in DCM was then added over 10 min. The reaction was stirred at RT until completion. The resulting solution was diluted with $H_2O$ (20 mL), extracted with DCM (3 × 20 mL), dried over $MgSO_4$ and concentrated *in vacuo*. The crude residue was purified by column chromatography (33% Pet. Ether in $Et_2O$) to afford the title compound, as a white foam (43 mg, 90%).

$^1$H NMR (400 MHz, Methanol-$d_4$) δ 1.92 – 2.01 (m, 1H), 2.30 – 2.38 (m, 1H), 3.10 – 3.31 (m, 2H), 4.19 – 4.28 (m, 2H), 6.70 (d, *J* = 8.7 Hz, 1H), 7.25 (dd, *J* = 8.8, 2.5 Hz, 1H), 7.45 – 7.51 (m, 2H), 7.78 (dt, 1H), 7.82 (dt, *J* = 7.9, 1.3, 1.3 Hz, 1H), 8.01 (t, *J* = 1.9, 1.9 Hz, 1H). $^{13}$C NMR (101 MHz, Methanol-$d_4$) δ 32.02, 51.07, 63.16, 67.31, 111.93, 118.46, 122.47, 125.24, 128.23, 129.23, 129.65, 130.66, 131.68, 135.15, 142.84, 153.83. MS: *m/z* (ES) 474 $[M-H]^-$. HRMS, found 473.9025 ($C_{16}H_{14}NO_4SBr_2$, $[M-H]^-$, requires 473.9010). TLC (hexane:EtOAc - 1:1) $R_f$: 0.43.

#### SYNTHESIS OF ARYL AZIDE PROBE 2

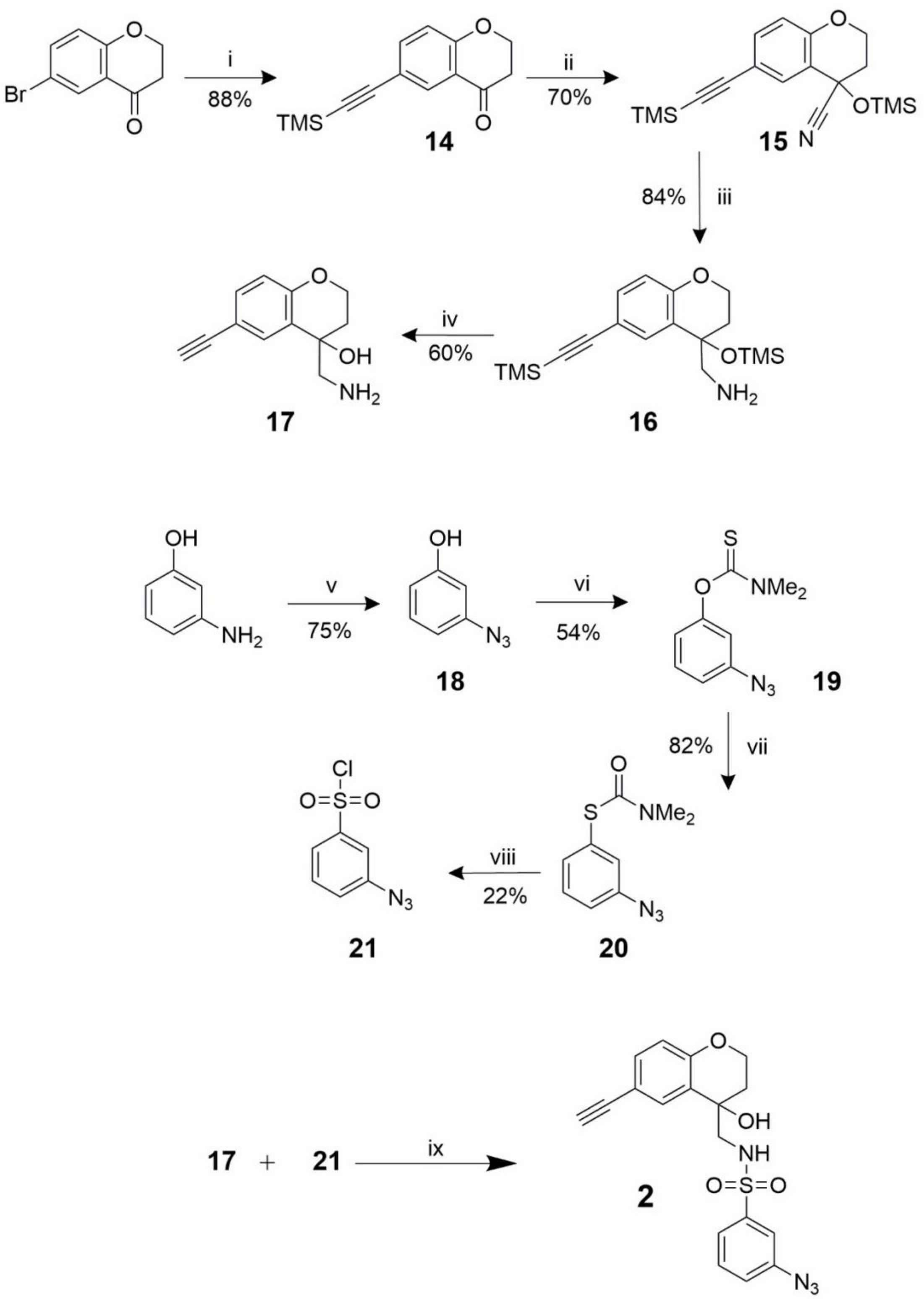


**SCHEME 2: Synthetic route to aryl azide containing probe, probe 2.** **i** - CuI, $PdCl_2(PPh_3)_2$, TEA, $C_5H_{10}Si$, DCM, 70°C, 24 h. **ii** - $ZnI_2$, TMS-CN, 50°C, ON. **iii** - $LiAlH_4$, THF, 0°C, 3 h. **iv** - $K_2CO_3$, DCM:MeOH (5:2), RT, 5 h. **v** - HCl/$H_2O$, $NaNO_2$, 0°C, 10 min. Followed by $NaN_3$, 0°C, 1 h. **vi** - DABCO, NMP, 50°C, 24 h. **vii** - Pd(t-$Bu_3$P)$_2$, Toluene, 100°C, 72 h. **viii** - NCS, HCl, ACN, 0°C, 6 h. **ix** – TEA, DCM, RT, 5 h.

**Compound 14** - 6-((trimethylsilyl)ethynyl)chroman-4-one

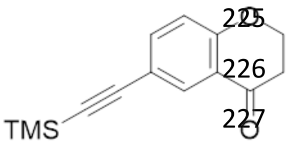


A solution of 6-bromo-4-chromanone (1.2 g, 5.3 mmol), CuI (30 mg, 0.2 mmol) and TEA (18 mL, 15.9 mmol) was sparged with argon. Bis(triphenyl phosphine) palladium chloride (112 mg, 0.2 mmol) and trimethylsilylacetylene (2.2 mL, 15.9 mmol) were added, the mixture was heated to 75 °C and stirred for 24 h. The resulting solution was diluted into $Et_2O$ (20 mL), washed with $NH_4Cl$:brine 9:1 (3 × 20 mL), washed with brine (60 mL), dried over $MgSO_4$ and concentrated *in vacuo*. The crude residue was purified by column chromatography (0 to 50 % EtOAc in Hexane) to afford the title compound, as a pale yellow oil (1.1 g, 88%).

$^1$H NMR (400 MHz, $CDCl_3$) δ 8.01 (d, J = 2.0 Hz, 1H, ArCH), 7.53 (dd, J = 8.6, 2.2 Hz, 1H, ArCH), 6.90 (d, J = 8. Hz, 1H, ArCH), 4.58 – 4.50 (m, 2H, $OCH_2CH_2$), 2.85 – 2.77 (m, 2H, $OCH_2CH_2$), 0.23 (s, 9H, TMS).$^{13}$C NMR (101 MHz, $CDCl_3$) δ 191.0, 161.7, 139.0, 131.2, 121.1, 118.2, 116.7, 103.8, 94.0, 67.2, 37.7, 0.1.

**Compound 15** - 6-((trimethylsilyl)ethynyl)-4-((trimethylsilyl)oxy)chromane-4-carbonitrile

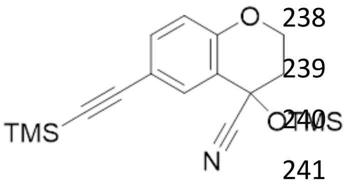


Compound **14** (730 mg, 3.0 mmol) and $ZnI_2$ (19 mg, 0.06 mmol) were suspended in DCM and cooled to 0 °C. Trimethylsilyl cyanide (560 µL, 4.5 mmol) was added dropwise and the resulting solution was stirred overnight. The resulting solution was diluted with DCM (20 mL), washed with $NaHCO_3$ (3 × 20 mL), extracted with DCM (2 × 60 mL), dried over $MgSO_4$ and concentrated *in vacuo* to afford the title compound, as an orange oil (718 mg, 70%). Due to column instability, compound **15** was carried forward without further purification.

$^1$H NMR (400 MHz, Methanol-$d_4$) δ 7.58 (d, J = 2.0 Hz, 1H, ArCH, 7.37 (dd, J = 8.6, 2.1 Hz, 1H, ArCH), 6.83 (d, J = 8.6 Hz, 1H, ArCH), 4.43 – 4.34 (m, 2H, $OCH_2CH_2$), 2.53 – 2.35 (m, 2H, $OCH_2CH_2$), 0.26 (s, 9H, TMS), 0.21 (s, 9H, TMS).

**Compound 16** - (6-((trimethylsilyl)ethynyl)-4-((trimethylsilyl)oxy)chroman-4-yl)methanamine

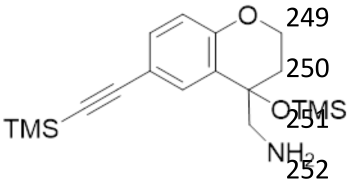


Compound **15** (718 mg, 2.1 mmol) was dissolved in THF and added drop-wise, over 15 min, to a stirred suspension of $LiAlH_4$ (319 mg, 8.4 mmol) in THF pre-cooled to 0°C. The reaction was stirred at 0°C until completion. The reaction was quenched using the Fieser procedure, the following solutions were added slowly at 0 °C: dilute with $Et_2O$, $H_2O$ (1 × w/v mass of $LiAlH_4$), 15% NaOH (1 × w/v mass of $LiAlH_4$), $H_2O$ (1.5 × w/v mass of $LiAlH_4$). The solution was warmed to RT, stirred for 15 min, dried over $MgSO_4$, filtered and concentrated *in vacuo* to afford the title compound, as an orange oil (425 mg, 84%).

$^1$H NMR (400 MHz, Methanol-$d_4$) δ 7.46 (d, J = 2.1 Hz, 1H, ArCH), 7.24 (dd, J = 8.5, 2.1 Hz, 1H, ArCH), 6.74 (d, J = 8.5 Hz, 1H, ArCH), 4.35 – 4.26 (m, 1H, $COHCH_2NH_2$), 4.24 – 4.17 (m, 1H,

$COHCH_2NH_2$), 3.00 – 2.86 (m, 2H, $OCH_2CH_2$), 2.30 – 2.17 (m, 1H, $OCH_2CH_2$), 2.14 – 2.03 (m, 1H, $OCH_2CH_2$), 0.22 (s, 9H, TMS), -0.02 (s, 9H, TMS).

**Compound 17** - 4-(aminomethyl)-6-ethynylchroman-4-ol

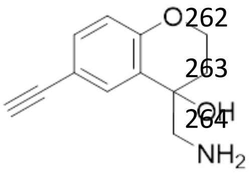


Compound **16** was shown by $^1$H NMR to be a mixture of trimethylsilyl (TMS) protected/ deprotected compound, therefore was fully deprotected before further purification.

Compound **16** (425 mg, 1.2 mmol) was dissolved in DCM in MeOH (5:2, 5 mL), before $K_2CO_3$ (200 mg, 1.5 mmol) was added and solution stirred at RT for 5 h. The resulting solution was diluted with $H_2O$ (20 mL), extracted with DCM (3 × 20 mL), washed with $NaHCO_3$ (2 × 60 mL), dried over $MgSO_4$ and concentrated *in vacuo*. The crude residue was purified by column chromatography (2% to 5% 7M $NH_3$ MeOH in DCM) to afford the title compound, as an orange oil (150 mg, 60%).

$^1$H NMR (400 MHz, Methanol-$d_4$) δ 7.56 (d, J = 2.1 Hz, 1H, ArCH), 7.24 (dd, J = 8.5, 2.1 Hz, 1H, ArCH), 6.74 (d, J = 8.5 Hz, 1H, ArCH), 4.29 – 4.18 (m, 2H, $COHCH_2NH_2$), 2.97 – 2.82 (m, 2H, $OCH_2CH_2$), 2.24 – 2.15 (m, 1H, $OCH_2CH_2$), 2.00 – 1.91 (m, 1H, $OCH_2CH_2$). $^{13}$C NMR (101 MHz, Methanol-$d_4$) δ 156.5, 133.6, 132.0, 128.5, 118.1, 115.5, 84.5, 76.9, 69.1, 64.6, 51.5, 33.2. MS: *m/z* (ES) 186 (100%, $[M-OH]^+$), 204 (85%, $[M+H]^+$). HRMS, found 204.1025 ($C_{12}H_{14}NO_2$, $[M+H]^+$, requires 204.1025). TLC (DCM:MeOH – 90:10) $R_f$: 0.14.

**Compound 18 - 3-azidophenol**

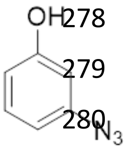


3-Aminophenol (400 mg, 3.6 mmol) was dissolved in $H_2O$ (1.6 mL) and HCl (1.6 mL) and cooled to 0 °C. $NaNO_2$ was dissolved in $H_2O$ (10 mL), cooled to 0 °C and added dropwise to the reaction over 10 min. The reaction was stirred 20 min at 0°C, before $NaN_3$ (240 mg, 3.6 mmol) in $H_2O$ (4 mL) was added dropwise over 10 min. The reaction was stirred for 90 min at 0 °C, before dilution in $H_2O$ (20 mL). The aqueous phase was extracted with $Et_2O$ (3 × 30 mL) and the combined organic phases were washed with $NaHCO_3$ (3 × 100 mL), brine (2 × 100 mL), dried over $MgSO_4$ and concentrated *in vacuo*. The crude residue was purified by column chromatography (10% EtOAc in Hexane) to afford the title compound, as a colourless oil (400 mg, 75%).

$^1$H NMR (400 MHz, $CDCl_3$) *δ* 7.21 (t, J = 8.1 Hz, 1H, ArC**H**), 6.68 – 6.59 (m, 2H, ArC**H**), 6.52 (t, J = 2.2 Hz, 1H, ArC**H**), 5.87 (s, 1H, ArCO**H**). $^{13}$C NMR (101 MHz, $CDCl_3$) *δ* 156.7, 141.5, 130.9, 112.3, 111.7, 106.4. Analysed by analytical LC-MS 20-98% ACN in $H_2O$ $R_t$ = 8.65 min, $ES^-$ $(M - H)^-$ m/z = 134.

**Compound 19 - O-(3-azidophenyl) dimethylcarbamothioate**

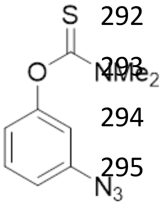


Compound **18** (400 mg, 2.9 mmol) and 1,4-diazabicyclo[2.2.2]octane (395 mg, 3.2 mmol) were dissolved in NMP (15 mL), before dimethyl thiocarbamoyl chloride (406 mg, 3.6. mmol) was added portion-wise over 10 min. The reaction was stirred at 50°C for 48 h and the resulting solution was diluted EtOAc (50 mL), washed with $H_2O$ (3 × 50 mL), brine (2 × 50 mL), dried over $MgSO_4$ and concentrated *in vacuo*. The crude residue was purified by column chromatography (8 to 15% EtOAc in Hexane) to afford the title compound, as a colourless oil (354 mg, 54%).

$^1$H NMR (400 MHz, $CDCl_3$) $\delta$ 7.34 (t, J = 8.1 Hz, 1H, ArC**H**), 6.91 (ddd, J = 8.1, 2.2, 0.9 Hz, 1H, ArC**H**), 6.86 (ddd, J = 8.2, 2.2, 0.9 Hz, 1H, ArC**H**), 6.76 (t, J = 2.2 Hz, 1H, ArC**H**), 3.43 (s, 3H, $N(CH_3)_2$), 3.32 (s, 3H, $N(CH_3)_2$). $^{13}$C NMR (101 MHz, $CDCl_3$) $\delta$ 187.2, 154.9, 140.9, 130.0, 119.5, 116.5, 113.9, 43.3, 38.7.

**Compound 20 - S-(3-azidophenyl) dimethylcarbamothioate**

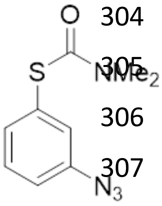


Compound **19** (100 mg, 0.45 mmol) was dissolved in toluene (5 mL) and heated to 100°C under argon. $Pd(tBu_3P)_2$ (5.0 mg, 2 mol%) was added and the reaction stirred 72 h at 100°C. The resulting solution was concentrated *in vacuo* and the crude residue was purified by column chromatography (5 to 20 % EtOAc in Hexane) to afford the title compound, as a colourless oil (82 mg, 82%). Reaction was monitored by IR, for the appearance of an indicative carbonyl stretch at 1667.13 $cm^{-1}$.

$^1$H NMR (400 MHz, $CDCl_3$) $\delta$ 7.38 (t, J = 7.8 Hz, 1H, ArC**H**), 7.32 – 7.28 (m, 1H, ArC**H**), 7.22 (t, J = 2.0 Hz, 1H, ArC**H**), 7.09 – 7.04 (m, 1H, ArC**H**), 3.15 – 3.01 (m, 6H, $N(CH_3)_2$). $^{13}$C NMR (101 MHz, $CDCl_3$) $\delta$ 166.2, 140.6, 132.2, 130.1, 126.1, 120.0, 37.0. IR vmax (neat, $cm^{-1}$) 2900 ($N_3$), 1667 (CO). HRMS, found 223.0650 ($C_9H_{114}OS$, $[M + H]^+$, requires 223.0654).

A small amount of starting material was observed in the $^1$H and $^{13}$C spectra, the crude mixture was carried through to next reaction.

**Compound 21 - 3-azidobenzenesulfonyl chloride**

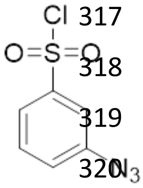


Compound **20** (85 mg, 0.38 mmol) was suspended in ACN (2.6 mL)/HCl (2M, 530 *m*L) and cooled to 0 °C. N-chlorosuccinimide (200 mg, 1.5 mmol) was dissolved in ACN (4 mL) and added dropwise to the reaction. The resulting solution was stirred at 0°C for 6 h and diluted with $iPr_2O$ (20 mL). The organic phase was washed with $H_2O$ (3 × 30 mL) and brine (2 × 30 mL), dried over $MgSO_4$ and concentrated *in vacuo*. The crude residue was purified by column chromatography (5 to 30% EtOAc in Hexane) to afford the title compound, as a colourless oil (18 mg, 22%).

$^1$H NMR (400 MHz, $CDCl_3$) $\delta$ 7.83 – 7.78 (m, 1H, ArC**H**), 7.67 (t, J = 2.1 Hz, 1H, ArC**H**), 7.62 (t, J = 8.0 Hz, 1H, ArC**H**), 7.40 – 7.35 (m, 1H, ArC**H**). $^{13}$C NMR (101 MHz, $CDCl_3$) $\delta$ 145.8, 142.4, 131.3, 125.6, 123.1, 117.4.

**Compound 2 - 3-azido-N-((6-ethynyl-4-hydroxychroman-4-yl)methyl)benzenesulfonamide**

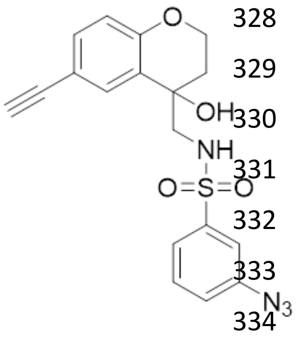


Compound **17** (22 mg, 0.078 mmol) was dissolved in DCM under an inert atmosphere at 0°C, before Triethylamine (TEA) (27 µL, 0.195 mmol) was added dropwise. A solution of compound **21** (20 mg, 0.094 mmol) in DCM was then added over 10 min. The reaction was stirred at RT until completion. The resulting solution was diluted with $H_2O$ (20 mL), extracted with DCM (3 × 20 mL), dried over $MgSO_4$ and concentrated *in vacuo*. The crude residue was purified by column chromatography (25% EtOAc in Hexane) to afford the title compound, as a white foam (15 mg, 56%). A 10 mM DMSO stock was prepared for biological testing.

$^1$H NMR (400 MHz, DMSO-$d_6$) $\delta$ 7.87 (s, 1H, CO**H**$CH_2$NH), 7.62 – 7.57 (m, 2H, ArC**H**), 7.50 (s, 1H, ArC**H**), 7.44 (d, J = 2.1 Hz, 1H, ArC**H**), 7.39 – 7.32 (m, 1H, ArC**H**), 7.23 (dd, J = 8.5, 2.0 Hz, 1H, ArC**H**), 6.74 (d, J = 8.4 Hz, 1H, ArC**H**), 5.46 (s, 1H, $SO_2$N**H**), 4.31 – 4.09 (m, 2H, OC$\mathbf{H}_2$$CH_2$), 3.98 (s, 1H, C≡C**H**), 3.12 (d, J = 13.3 Hz, 1H, COHC$\mathbf{H}_2$NH), 2.94 (d, J = 13.2 Hz, 1H, COHC$\mathbf{H}_2$NH), 2.29 – 2.16 (m, 1H, $OCH_2$C$\mathbf{H}_2$), 1.92 – 1.78 (m, 1H, $OCH_2$C$\mathbf{H}_2$). $^{13}$C NMR (101 MHz, DMSO-$d_6$) $\delta$ 154.7, 142.4, 140.6, 132.2, 131.4, 130.8, 127.2, 123.0, 122.9, 116.9, 116.8, 113.2, 83.7, 78.9, 66.6, 63.1, 50.8, 31.9. HRMS, found 383.0820 ($C_{18}H_{15}N_4O_4S$, $[M - H]^-$, requires 383.0814).

**$IC_{50}$ determination by male gamete formation assay**

The male gamete formation assay (MGFA) was adapted from the previously described dual gamete formation assay (DGFA)[3] to identify male gamete targeted activity. Briefly, compounds were dispensed into 384-well plates using the HP D300 Digital Dispenser in serial dilutions. Single point controls, a DMSO negative control (0.25% final assay volume) and 20 μM **DDD01028076** positive control, were included across the plate. Maintaining RT, 10 μl ookinete medium was dispensed into each well, followed by addition of 50 μl of stage V NF54 gametocyte culture (day 14 post-induction and onwards). Gametocyte cultures, demonstrating ≥0.2% exflagellation, were diluted to 25 million erythrocytes/ml for the assays. Plates were then immediately incubated at 4°C for 4 minutes, followed by 28°C for 5 minutes.

Exflagellation events were recorded using a Nikon Eclipse Ti inverted microscope by automated data capture. Using the JOBS module of NIS elements, a 10-frame time-lapse was captured at 4 frames per second under phase contrast at x4 objective, 1.5x zoom. Time-lapses were recorded at a z-coordinate determined from a prior autofocus step.

A semi-automated algorithm designed in Icy Bioimage Analysis software was used to obtain an exflagellation centre count based on the size, pixel intensity and circularity of disruptions in the erythrocyte monolayer of each time-lapse. Raw data with a Z' factor ≥0.4 was normalised to positive and negative controls to calculate a percentage inhibition using Eq. 1:

$$\%\,inhibition = 100 - \left(\left(\frac{test\ compound - positive\ control}{negative\ control - positive\ control}\right) \times 100\right)$$

The $IC_{50}$ of each compound was then determined using GraphPad Prism software.
